# CRISPR interference screens reveal tradeoffs between growth rate and robustness in *Synechocystis* sp. PCC 6803 across trophic conditions

**DOI:** 10.1101/2023.02.13.528328

**Authors:** Rui Miao, Michael Jahn, Kiyan Shabestary, Elton Paul Hudson

## Abstract

Barcoded mutant libraries are a powerful tool for elucidating gene function in microbes, particularly when screened in multiple growth conditions. Here, we screened a pooled CRISPR interference library of the model cyanobacterium *Synechocystis sp*. PCC 6803 in 11 bioreactor-controlled conditions, spanning multiple light regimes and carbon sources. This gene repression library contained 21,705 individual mutants with high redundancy over all open reading frames and non-coding RNAs. Comparison of the derived gene fitness scores revealed multiple instances of gene repression being beneficial in one condition while generally detrimental in others, particularly for genes within light harvesting and conversion, such as antennae components at high light and PSII subunits during photoheterotrophy. Suboptimal regulation of such genes likely represents a tradeoff of reduced growth speed for enhanced robustness to perturbation. The extensive dataset assigns condition-specific importance to many previously unannotated genes, and suggests new functions for central metabolic enzymes. Prk, GAPDH, and CP12 were critical for mixotrophy and photoheterotrophy, which implicates the ternary complex as important for redirecting metabolic flux in these conditions in addition to inactivation of the Calvin cycle in the dark. To predict the potency of sgRNA sequences, we applied machine learning on sgRNA sequences and gene repression data, which showed the importance of C enrichment and T depletion in the first 12 bp proximal to the PAM site. Fitness data for all genes in all conditions is compiled in an interactive web application.

## Introduction

Photoautotrophic microbes, including various cyanobacterial strains, have been pursued as next-generation catalysts for synthesis of fuels or chemicals (Lips et al. 2018). In addition to their biotechnological potential, cyanobacteria have served as model organisms for photosynthesis and carbon fixation, due to an ancestral relationship to the chloroplasts of photosynthetic eukaryotes. Nevertheless, 45% of genes in the most-studied cyanobacterium *Synechocystis* sp. PCC 6803 are not annotated with a function, compared to 15-35% in the model heterotroph *E. coli* (Ghatak et al. 2019). This knowledge gap is a significant obstacle to fully understanding cyanobacterial physiology, as well as to developing cyanobacteria into efficient cell factories for bioproduction. In recent years, there have been a number of technical advances that accelerate high-throughput functional genomics. Barcoded mutant libraries, created *via* transposon, CRISPR/Cas, or CRISPRi, allow mutant tracking via NGS, and thus the screening of thousands of genes simultaneously across environmental conditions (M. N. Price et al. 2018; T. Wang et al. 2018; Jahn et al. 2021; Yao et al. 2020; Garst et al. 2017; Vo et al. 2021). Such a highly parallel experimental format facilitates identification of condition-specific gene fitness.

Over the course of evolution, microorganisms have acquired the genetic inventory and regulatory mechanisms to provide quick responses toward dynamic environments. For example, the lithoautotroph *Cupriavidus necator* simultaneously expresses multiple alternative pathways for substrate assimilation (Jahn et al. 2021), and *Pseudomonas putida* precautionarily expresses numerous efflux pumps to increase solvent and xenobiotic tolerance (Ramos et al. 2015). A systems biology analysis suggested that *Synechocystis* does not efficiently regulate proteins involved in light harvesting and CO_2_ fixation in high-growth conditions, possibly retaining excess enzyme capacity in anticipation of changing conditions (Jahn et al. 2018), but the same expression can be burdensome for growth in another environment. Robustness in the presence of light fluctuations is a well-studied phenomenon in photosynthetic organisms. In plants, several non-photochemical quenching (NPQ) routes are sustained when conditions are no longer stressful due to low expression of key enzymes, so that rapid deactivation through additional enzyme expression is an objective for increasing crop yield (Kromdijk et al. 2016). Again, the high parallelization of barcoded library functional genomics allows the study of possible sub-optimal, or “wasteful”, gene expression across the genome and across multiple conditions.

Here, we have used a CRISPR interference library to elucidate the contribution of *Synechocystis* genes to cell growth in specific conditions and discover fitness trade-offs that highlight evolutionary pressures to adapt to changing environments. A previous CRISPRi library for *Synechocystis* contained two single guide RNAs (sgRNA) targeting each gene, which resulted in ambiguous results when one sgRNA gave a growth phenotype and the other did not (Yao et al. 2020). Our expanded CRISPRi library has the majority of genes targeted by five sgRNAs to increase confidence in derived fitness scores, as well as sgRNAs targeting non-coding RNAs (sRNAs, antisense RNAs, and alternative transcription start sites), a widespread class of potentially regulatory molecules in cyanobacteria (Kopf and Hess 2015). The expanded CRISPRi library was cultivated in 11 different conditions where carbon source, nitrogen source, and light availability varied. The resulting large datasets reveal previously unannotated genes as being important for cell growth in certain conditions, as well as several examples of growth-robustness tradeoffs, where gene repression accelerated growth in one condition but was detrimental in others. All analyzed fitness data for competition experiments can be accessed on an interactive web application (https://m-jahn.shinyapps.io/ShinyLib/).

## Results

### An expanded CRISPRi library for the cyanobacterium *Synechocystis* sp. PCC 6803

The CRISPRi library was constructed so that both the catalytically inactive Cas9 enzyme (dCas9) and sgRNA are transcribed by anhydrotetracycline (aTc)-inducible promoters. Upon addition of aTc, the dCas9 and sgRNA are expressed, and the dCas9-sgRNA complex mediates blockage of transcription, with gene specificity determined by the sgRNA sequence (Qi et al. 2013; Yao et al. 2016, 2020). Up to five sgRNAs were designed to target each gene and non-coding RNA (ncRNA) on the chromosome and the native plasmids of *Synechocystis*. Five sgRNAs were designed for most genes (92%), while fewer sgRNAs were generated for shorter ORFs and for ncRNAs (25% of ncRNAs were targeted by five sgRNAs) (Figure 1 A). The resulting 21,705 sgRNAs were synthesized as a pool (GenScript) and integrated into a *Synechocystis* strain harboring dCas9. All transformants, where each cell contained a single sgRNA, were scraped off agar plates and this constituted the pooled CRISPRi library (Methods). The presence of all designed sgRNAs in the pooled library was verified by NGS.

**Figure 1:**
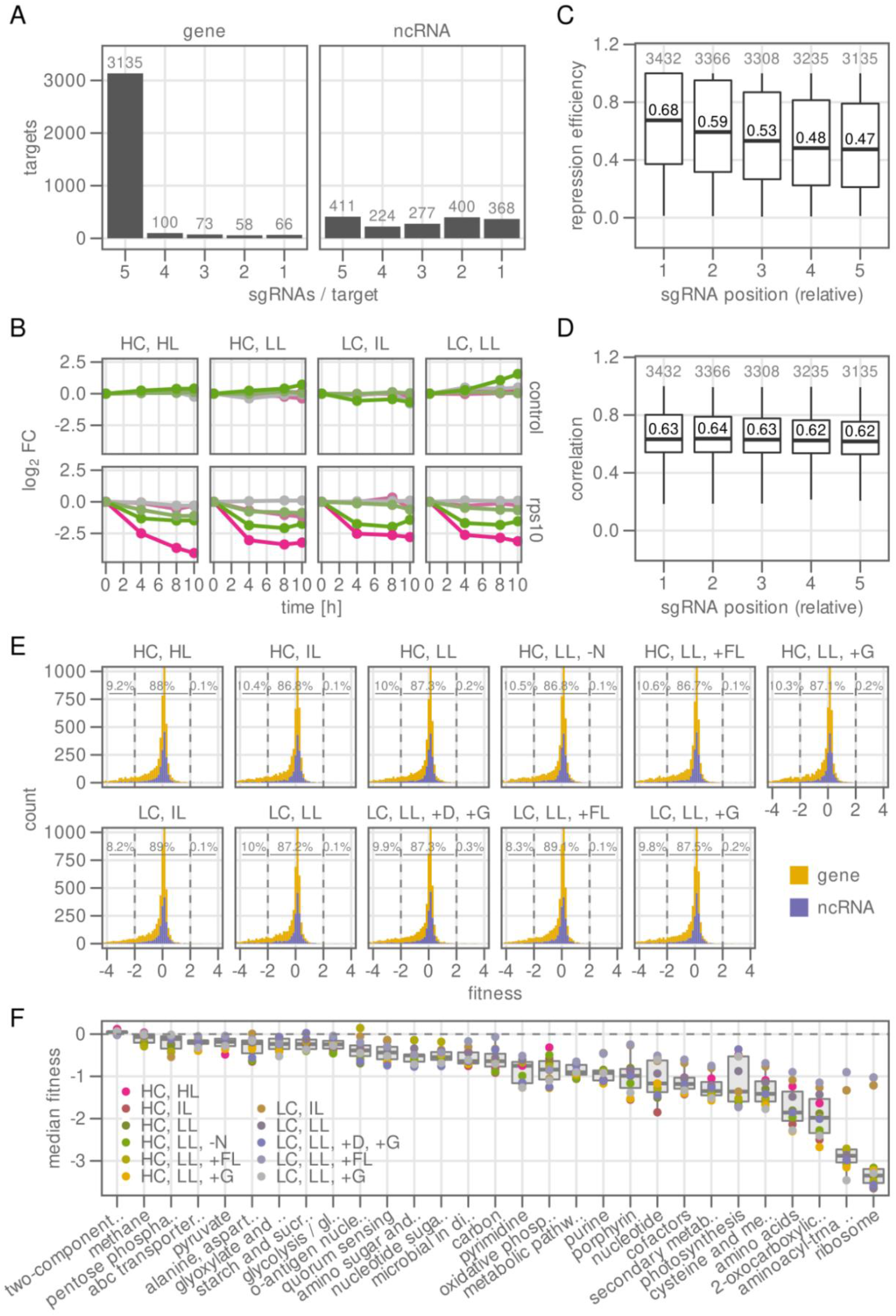
*Synechocystis* CRISPRi library targets most genes with 5 sgRNAs. A CRISPRi repression library was constructed for *Synechocystis* sp. PCC 6803 targeting 3,432 genes and 1,712 non-coding RNAs. **A)** Almost all genes (3136 of 3432, 91%) were targeted with 5 sgRNAs, while ncRNAs were often targeted with fewer sgRNAs because of limited sequence length. **B)** Abundance log_2_ fold-change of example mutants over a time course from 0 to 10 generations in 4 different conditions. Upper row: 5 non-targeting sgRNA controls, lower row: 5 sgRNAs targeting the ribosomal gene *rps10*. **C)** Relative repression efficiency of an sgRNA depending on distance to the start codon (1 - closest, 5 - most distant). The sgRNA with the strongest effect on fitness is set to a value of 1. Only sgRNAs targeting genes were included for the analysis. **D)** Pearson’s correlation coefficient of each sgRNA to the other sgRNAs targeting the same gene, depending on distance to the promoter. The correlation coefficient was rescaled to a range of 0 to 1. The same set of sgRNAs as in C) was used. **E)** Histogram of fitness score per condition, for genes (yellow) and ncRNAs (purple). Inset numbers show the percentage of genes falling in three different bins: strong negative fitness (−4 to -2), no or weak effect on fitness (−2 to 2), strong positive fitness (2 to 4). **F)** Median fitness score of genes per pathway (KEGG), broken down by cultivation condition (see Table 1).

We performed turbidostat cultivations of the pooled library with varying CO_2_ concentration, light exposure, and additional treatments such as the addition of glucose to enable mixotrophic growth, the addition of glucose and DCMU for photoheterotrophic growth, and nitrogen starvation (Table 1). Conditions were expected to put demands on carbon and energy metabolism, which cyanobacteria must balance to maximize growth rate. The composition of the pool in each condition was determined by NGS of sgRNA cassettes after 0, 4, 8 and 10 generations. The change in abundance of each library member over the cultivation duration is a measure of the relative growth rate of that member, and was used to determine a fitness score (Love, Huber, and Anders 2014; Yao et al. 2020) (Figure 1 B). Each sgRNA for a gene may have a different efficacy *in vivo*, and thus different magnitude of effect on cell growth (fitness score). We found that sgRNAs targeting most proximal to the promoter of the target gene (“position 1”) had the highest absolute fitness scores, and thus presumably higher repression efficiency (median efficiency = 0.67, where relative fitness is scaled between 0 and 1; Figure 1 C). The absolute fitness score of sgRNAs declined with distance from the promoter (position 5, median efficiency = 0.47), which is similar to previous reports (T. Wang et al. 2018). We also confirmed that the position of an sgRNA has no effect on the correlation with other sgRNAs for the same gene (Figure 1 D). Gene fitness scores were calculated from individual sgRNA fitness scores by weighted mean, and significance was determined by calculating the multiple hypothesis adjusted *p*-value (padj) from individual sgRNA fitness scores of the same target (Methods).

**Table 1.** Summary of the conditions used in turbidostat cultivation of the pooled library.

| Condition ID | CO<sub>2</sub> supplement | Light intensity (mmol m<sup>-2</sup> s<sup>-1</sup>) | Treatment |
| --- | --- | --- | --- |
| HC, HL | 1 % | 1000 | - |
| HC, IL | 1 % | 150 | - |
| HC, LL | 1 % | 60 | - |
| HC, LL, -N | 1 % | 60 | BG11<sub>0</sub> |
| HC, LL, +FL | 1 % | 60/1500 | 5 seconds of fluctuating light every 5 minutes |
| HC, LL, +G | 1 % | 60 | 5mM glucose |
| LC, IL | 0.04% | 150 |  |
| LC, LL | 0.04% | 60 |  |
| LC, LL, +FL | 0.04% | 60/1500 | 5 seconds of fluctuating light every 5 minutes |
| LC, LL, +G | 0.04% | 60 | 5mM glucose |
| LC, LL, +D, +G | 0.04% | 60 | 10mM DCMU and 5mM glucose |

The majority of targets that showed an effect on fitness were genes (Figure 1 E), while only few ncRNAs showed an independent effect upon repression. To obtain an overview of the pathways most important for cell fitness, genes were sorted by KEGG pathways and the median fitness per pathway in each condition was calculated (Figure 1 F). As expected, most genes that show an effect on fitness tend to show a reduction in fitness, resulting in negative median fitness scores for most pathways in different conditions. The metabolic pathways showing the strongest detrimental effect on fitness upon repression were associated with ribosomes, nucleotide and tRNA biosynthesis, and amino acid biosynthesis. Non-central dogma pathways with strong average fitness loss include photosynthesis, biosynthesis of cofactors and secondary metabolites including chlorophyll, and oxidative phosphorylation. Interestingly, all of these pathways were associated with energy metabolism, while the carbon metabolic pathways showed milder effects, with first appearances for central carbon metabolism and glycolysis and gluconeogenesis at rank 15 and 21, repsectively. This relatively weak fitness effect could be due to high enzyme abundance in central carbon metabolism (underutilization of enzymes), a higher fraction of isoenzymes than in energy metabolism, or compensation by redirection of metabolic flux through other pathways. In two environmental conditions, low CO_2_ with intermediate light (LC,IL) and low CO_2_ with fluctuating light (LC,LL,+FL), the fitness penalty for repression of ribosomal and tRNA biosynthesis genes was significantly weaker than in other conditions, while other essential genes, such as photosynthesis, had fitness penalties similar to other conditions (Figure 1 F). Cells in these conditions were under extreme light stress and showed reduced growth rate, so that the demand for ribosomes was reduced. This reduced demand may result in a relative insensitivity to ribosome repression by CRISPRi compared to other conditions where cells grow faster.

In order to identify genes particularly important for fitness in changing carbon or light conditions, but not both, we applied multiple linear regression modeling to our fitness data. These models can dissect the influence of the single variables carbon concentration, light intensity and additional treatments on the fitness score of each gene. We selected 187 genes with significantly changed fitness in any condition (threshold: 4.0), clustered these using t-SNE, and overlaid the results from multiple linear regression (Figure S1). We found several genes (and gene clusters) of interest which we analyze in detail in the following sections, for example photosystem and phycobilisome subunits (*psb, apc*) in mixo- and photoheterotrophic conditions, Flv proteins (*sll0217, sll0219*) in fluctuating light, and the proposed CBB cycle regulator CP12 (*ssl3364*).

### Fitness tradeoffs for anticipating light stress and carbon limitation

Repression of the PSI subunits A, B, C, and D was detrimental for growth in all conditions, which is in agreement with their integral role in PSI formation and function (Yu et al. 1995; Malavath et al. 2018). Repression of the small PSI subunits E,F,J,K,L,I, and M also affected growth negatively, but in most conditions the effect was weak (Figure S2). This is in agreement with previous reports that the small subunits could be deleted in *Synechocystis* without effect on growth (Malavath et al. 2018; Jeanjean et al. 2008). Exceptions were conditions with low CO_2_ with intermediate or fluctuating light (LC,IL and LC,FL). The small PSI subunits may thus play a role in PSI function under light-stress conditions. PsaL and PsaI were previously shown to be critical for formation of the PSI trimer in *Synechocystis* (Netzer-El, Caspy, and Nelson 2018; Çoruh et al. 2021), though the physiological relevance of the PSI oligomerization is not fully elucidated. Repression of PSII reaction center proteins CP43 (*psbC*), D2 (*psbD*), and PsbK resulted in strong negative fitness effects (Figure 2 A). The D1 subunit of the reaction center is encoded by three genes, of which *psbA1* is often considered a neutral site and was used here to integrate the dCas9 enzyme. The repression of either of the two other copies, *psbA2* and *psbA3*, did not result in significant fitness change, most likely because they are 99% identical and compensate each other. When cells were grown on glucose in the presence of DCMU to block PSII activity (photoheterotrophy), most PSII genes showed growth advantage upon repression, suggesting that natural expression of PSII proteins are a fitness burden in this condition. Silencing of unused proteins may allow cells to reinvest resources into other functions (Jahn et al. 2018, 2021). PsbK was an exception, as it was also critical for cell growth even in photoheterotrophic conditions, in contrast to the pattern of the other PSII subunits.

**Figure 2:**
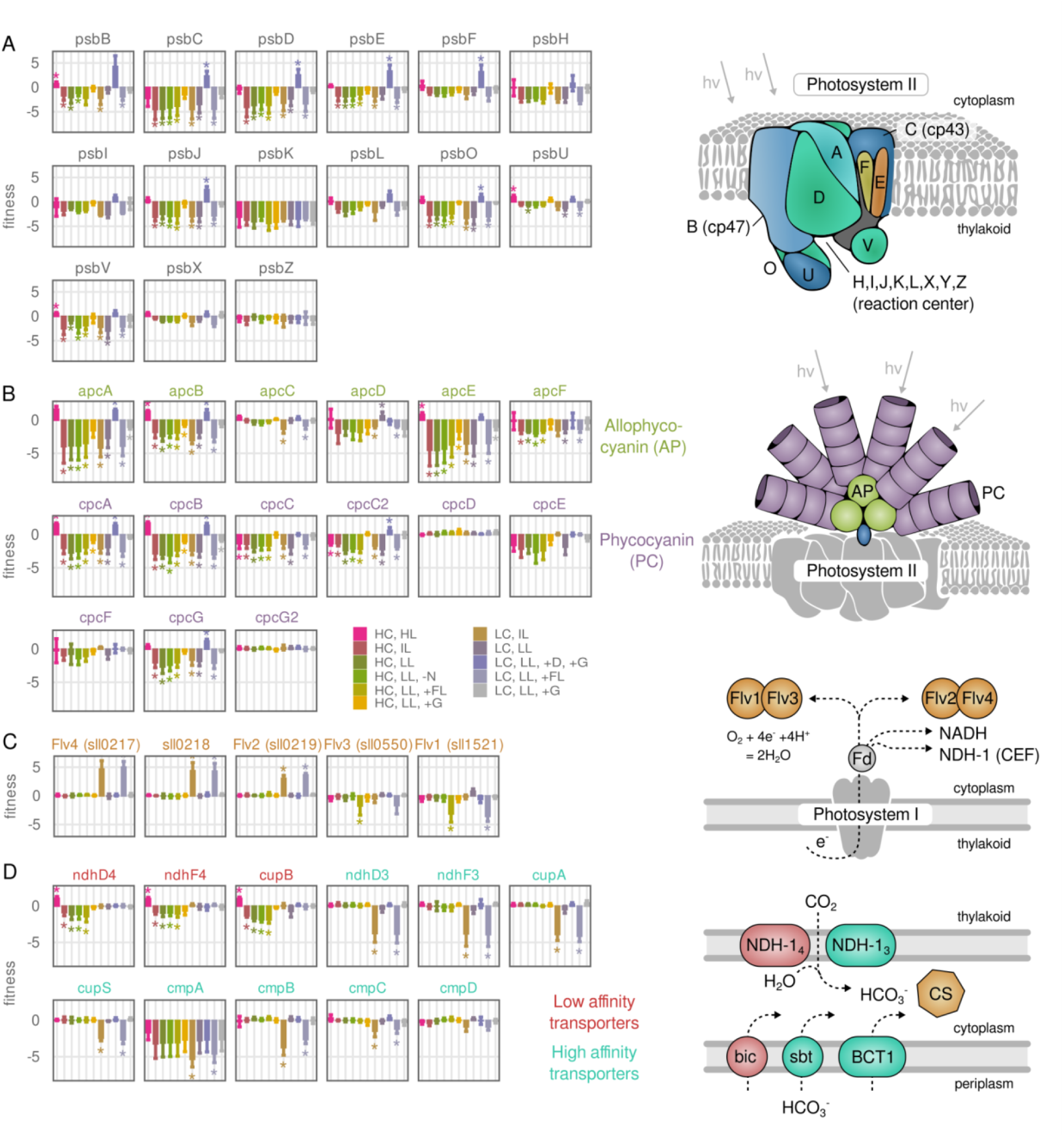
Adaptations to excess carbon or light are suboptimal in some environments. **A)** Fitness score for repression of selected genes encoding photosystem II subunits. Fitness score for all PSI and PSII subunits can be found in Figure S1. Asterisk: Wilcoxon rank sum test adjusted p-value ≤ 0.01. Illustration: Photosystem II structure adapted from KEGG (Kanehisa et al. 2017). **B)** Fitness score for repression of genes encoding phycobilisome subunits. Illustration: Phycobilisome structure adapted from KEGG **C)** Fitness score for repression of genes encoding flavodiiron (Flv) proteins. **D)** Fitness score for repressing genes encoding selected carbon transporters in *Synechocystis*: NDH-14 complex (*ndhD4, ndhF4, cupB*), NDH-1_3_ complex (*ndhD3, ndhF3, cupA, cupS*), BCT1 (*cmpABCD*). Red: low Ci affinity transporters, turquoise: high Ci affinity transporters.

The apparent wasteful expression of PSII in photoheterotrophic conditions motivated us to look for other instances of potential suboptimal regulation, particularly in light harvesting and conversion, where preparedness for a potential shift in light intensity could come at the expense of growth rate. One example is the expression of the phycobilisome (PBS), a large membrane-extrinsic protein complex serving as a photon-capturing antenna (Adir, Bar-Zvi, and Harris 2020). PBS is a significant proportion of the cellular proteome (20 to 40% by mass; Figure S3) (Jahn et al. 2018; Grossman et al. 1993), and consists of stacked rods of allophycocyanin (*apc* operon) and phycocyanin (*cpc* operon) subunits which each absorb light energy using phycobilin pigments (Arteni, Ajlani, and Boekema 2009). The expression of PBS genes is regulated in response to perceived light, with expression increasing at low light or in presence of DCMU and decreasing in high light (MacKenzie et al. 2005; Jahn et al. 2018; Zavřel et al. 2019). However, the cyanobacterium *Synechocystis* does not fully repress phycobilisomes even at extreme light, though photosynthesis can still occur when the entire PBS is knocked out (Ajlani et al. 1995). Instead, excessive PBS excitation is quenched in high light by the orange carotenoid protein (Domínguez-Martín et al. 2022). Additionally, PBS may partially detach from PSII to prevent energy transfer (Calzadilla & Kirilowsky, 2020). Several studies have proposed artificial antennae truncation to reduce the effective cross section for light capture and thus prevent wasteful NPQ in cyanobacteria cultures (Melis 2009; Grossman et al. 1993; Kirst, Formighieri, and Melis 2014; Lea-Smith et al. 2014). The CRISPRi fitness data showed a general negative effect of PBS repression across most conditions, except for extreme light (HC, HL) and photoheterotrophy (LC,LL,+G,+ D) conditions, where repression of ApcAB and CpcAB clearly improved growth rate (Figure 2 B, Figure S3). The magnitude of the growth defect of PBS repression was also reduced in the mixotrophy (+G) conditions. Thus, in most conditions repression of the antennae is detrimental, though in conditions where the antennae are not needed, repression led to an increase in growth rate (see Discussion for possible explanations).

A second example of a trade-off between robustness and growth rate was observed for the flavodiiron (Flv) proteins. Flvs protect cyanobacteria from excess light energy and thus over-reduction of NAD(P) by reducing molecular oxygen to water (Figure 2 C) (Allahverdiyeva et al. 2013). In *Synechocystis*, four Flv proteins are known (Flv1 to 4), of which Flv1-3 and Flv2-4 associate as heterodimers. Flv1-3 is constitutively expressed, essential for survival on intense fluctuating light, and caps excess light energy at low and high carbon levels (Santana-Sanchez et al. 2019). In agreement to this, we observed a significant negative effect on cell growth among Flv1 or Flv3 repression clones in fluctuating light (+FL) conditions regardless of CO_2_ concentration (Figure 2 C). Flv2 and Flv4 were reported to be involved in dissipating electron pressure from downstream of PSI *via* the Mehler-like reaction at constant light (Santana-Sanchez et al. 2019). Additionally, *sll0218*, a gene of unknown function encoded within the Flv2-Flv4 operon, was reported to play an important role in PSII assembly and stabilization (Bersanini et al. 2017). In the CRISPRi library, Flv2-4 repression clones had fitness increases that were specific to the two low Ci and excess light conditions (LC,IL and LC,LL,+FL). Two possible robustness-growth trade-offs could be the cause: one related to excess dissipation of electrons, and another related to protein burden. In excess light and low Ci conditions, Flv2-4 and one of the NAD(P)H dehydrogenase involved complexes - NDH-1_3_ are upregulated, presumably to accelerate electron dissipation and cyclic electron transfer around PSI (Santana-Sanchez et al. 2019; C. Zhang et al. 2020), which may lead to an excessive drain of electrons. On the other hand, O_2_ consumption by Flv2-4 was diminished at pH >7 (Santana-Sanchez et al. 2019). As the pH of the turbidostat cultures was maintained at 7.8, Flv2-4 may be partially inactive and instead constitute a protein burden. Thus, the increased fitness may be a consequence of saved resources from repression of un-utilized Flv2-4 – particularly as Flv2-4 transcription is upregulated in low CO_2_ conditions (P. Zhang et al. 2009).

Another adaptation to Ci availability is the regulation of Ci transporters. *Synechocystis* encodes five major Ci transport proteins (reviewed in (G. D. Price 2011)): two low affinity/high flux transporters, BicA *(bicA*) and NDH-1_4_ (*ndhD4, ndhF4, cupB*), and three high affinity/low flux transporters, SbtAB *(sbtA*), BCT1 (*cmpABCD*), and NDH-1_3_ (*ndhD3, ndhF3, cupA, cupS*). NDH-1_4_ is constitutively expressed for basal Ci uptake; expression of the other Ci transporters is induced in Ci-limited conditions (Shibata et al. 2001). The NDH-1_3/4_ complexes hydrate CO_2_ to HCO3-(+H^+^) on the cytoplasmic side, with the help of CO_2_ hydration proteins CupA, CupS, or CupB, while the others transport HCO3-across the cytoplasmic membrane. CRISPRi repression of the BicA and SbtAB transporters did not affect growth rate in any tested condition (Figure S4). In contrast, repressing the subunits of the low affinity transporter NDH-1_4_ reduced growth in most high carbon conditions and had no effect in low carbon conditions (Figure 2 D), indicating that NDH-1_4_ is the major Ci transporter when CO_2_ is abundant. However, repression of NDH-1_4_ increased growth in the high carbon and high light condition (HC,HL). It has been shown that, besides CO_2_ transportation, the NDH-1_3_ complex also plays an important role in PSI-mediated CEF, in a way that is distinct from the NDH-1L complex (Bernát et al. 2011). High light induces the expression of the NDH-1_3_ complex to accelerate cyclic electron flow (C. Zhang et al. 2020), as well as its high affinity CO_2_ transportation capacity. The additional presence of the low affinity transporter complex NDH-1_4_ could be a burden in the ‘HC, HL’ condition. In Ci limiting conditions, the high affinity transporters are essential for efficient carbon uptake, thus NDH-1_3_ and BCT1 repressing mutants showed significantly reduced fitness in two conditions characterized by strong imbalance of light and CO_2_ supply (LC,IL and LC,LL,+FL). Furthermore, the repression of genes involved in the C_i_ regulatory network, *ccmR* (also named *ndhR, rbcR*), *cmpR*, and *cyabrB2* (*sll0822*), also showed strong effects on cell fitness. For example, repression of low affinity C_i_ transporter inhibitor *ccmR* increased cell fitness in the LC, IL condition, and being consistent with previous studies, the repression of *cyabrB1* is lethal and the repression of *cyabrB2* also reduced cell fitness in all conditions (Ishii and Hihara 2008; Kaniya et al. 2013; Orf et al. 2016)(Figure S4).

### Essentiality of genes in central carbon metabolism

*Synechocystis*, like many other bacteria, has gene duplications or isoenzymes that can theoretically compensate for the loss or repression of a central carbon metabolism gene. Although the metabolic flux through most major reactions of central metabolism is known (Nakajima et al. 2014; You, He, and Tang 2015), it is often not easy to determine which genes/isoenzymes contribute most to carrying a reaction’s flux. The CRISPRi library accelerates the study of central metabolism by simultaneously screening all mutants with repression of a single gene in growth competition, as they are often laboriously studied with knockouts of single genes. It also has the advantage of targeting otherwise essential genes and thus gauging their relative importance (Donati et al. 2021). An essential reaction catalyzed by a single enzyme will lead to a strong fitness penalty when the corresponding gene is repressed. On the other hand, a reaction catalyzed by two isoenzymes will show either a partial fitness penalty (reduced flux) or no penalty at all (unchanged flux) if one enzyme can compensate for the loss of the other. We compared the fitness of central carbon metabolism genes in three growth conditions, phototrophy, mixotrophy and photoheterotrophy (Figure 3 A). Incorporation of glucose, which serves as either an additional or the sole source of carbon and energy in mixotrophy or photoheterotrophy, respectively, results in a significant change in metabolic flux patterns (Figure S5) (Nakajima et al. 2014). In cases where multiple genes are annotated for a reaction, comparison of fitness scores can reveal that one gene is more important than another. For example, we confirmed that *glk* (*sll0593*) but not *xylR* (*slr0329*) encodes the main hexokinase (HEX) for glucose import (Figure 3 A), which is consistent with a previous study comparing *glk/xylR* knockouts (Lee et al. 2005). Similarly, *cbbA* (*sll0018*) but not *fda* (*slr0943*) was identified to be responsible for around 90% of total fructose bisphosphate aldolase (FBA) activity (Nakahara et al. 2003), and we indeed observed reduced fitness scores in all photoautotrophic and mixotrophic conditions for *cbbA* repression (Figure 3 A). Furthermore, our results indicate that one of the pyruvate kinase (PYK) isoforms encoded by *sll1275* is more important in all conditions than the isoform encoded by *sll0587* (Figure 3 A), which was predicted to be highly inhibited by ATP in an *in silico* study (Haghighi 2021).

**Figure 3.**
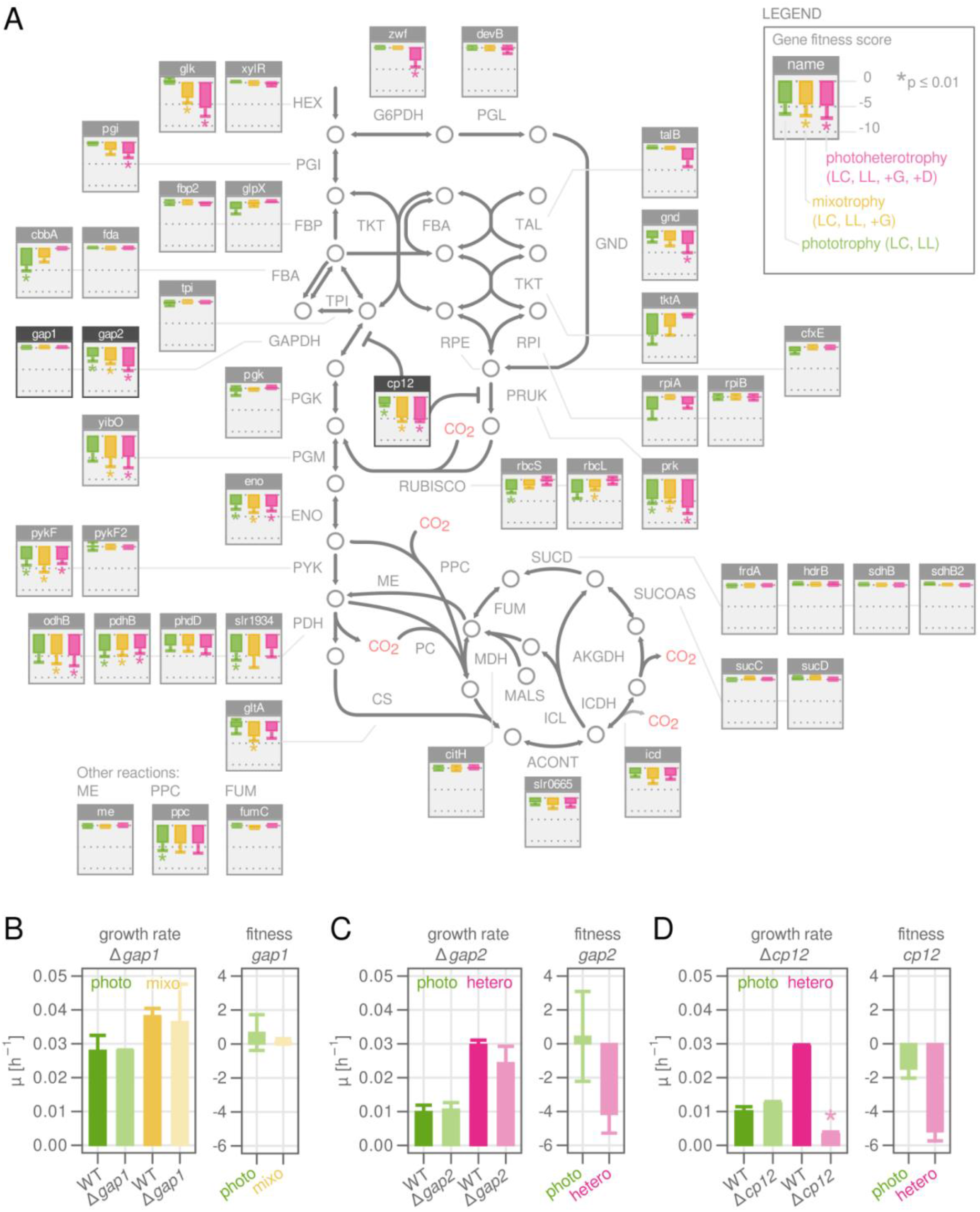
Essentiality of genes in central carbon metabolism. **A)** Metabolic map of central carbon metabolism for *Synechocystis sp*. PCC 6803. Reactions and their directionality are shown with arrows and named with capital letters according to the BiGG standard. Metabolites are shown as circles. Minifigures show fitness scores for the genes associated with the respective metabolic reaction. Green bars: phototrophy, yellow bars: mixotrophy, red bars: photoheterotrophy. Asterisk: p-value from Wilcoxon rank sum test lower than 0.01. **B)** Validation of CRISPRi library results for Gap1 using a Δ*gap1* deletion strain. Cultivation was performed in the same conditions as were used for cultivating the CRISPRi library, but in batch mode instead of turbidostat. Detailed growth curves are shown in Figure S5. Phototrophy: HC, LL. Mixotrophy: HC, LL, +G. WT was used as control in each cultivation condition, and the specific growth rate *μ* was calculated as the slope of log OD720 between hour 20 and 40. The fitness score of the corresponding gene is shown for comparison. Error bars: Mean and standard deviation of at least two biological replicates. Asterisk: p-value from Student’s t-test lower than 0.01. **C)** As in B) but for Gap2. Phototrophy: LC, IL. Photoheterotrophy: LC, LL, +G, +D. **D)** As in B) but for CP12. Phototrophy: LC, LL. Photoheterotrophy: LC, LL, +G, +D.

In general, gene fitness scores correlated well with expected enzyme usage based on reported fluxes (Figure S4). A high fitness penalty for the repression of the oxidative PPP (G6PDH, PGL, GND) was evident for photoheterotrophic growth, but not in the other conditions. The TCA cycle on the other hand carried very low flux in all conditions (Figure S4) and accordingly, repression of TCA associated genes had nearly no effect on fitness (Figure 3 A). It is important to keep in mind that enzyme abundance is usually higher than what is minimally required to maintain flux. Such a ‘reserve flux capacity’ was shown to increase robustness to perturbations or environmental changes in different bacteria (O’Brien et al., 2016, Mori et al., 2017, Sander et al., 2019). As a consequence, incomplete repression can leave sufficient residual enzyme capacity to maintain the flux of a reaction, which may complicate the interpretation of fitness scores. An example of this is Rubisco, which showed moderately but not dramatically reduced fitness for both subunits (*rbcS, rbcL*), despite being the primary source of carbon fixation during photoautotrophic growth. Another example is phosphoribulokinase (Prk), the committed step for CO_2_ fixation that provides the precursor ribulose-1,5-bisphosphate for Rubisco. Surprisingly, the growth penalty for Prk repression was highest for photoheterotrophy (Figure 3 A), a condition where flux through the enzyme has been shown to be low (Figure S4). The relative importance of Prk in photoheterotrophic conditions may be due to its role in the ternary complex Prk-CP12-Gap2.

We also found some discrepancies among the reported fluxes and calculated fitness scores for the associated enzymes. Phosphoglycerate kinase (PGK) and glyceraldehyde 3-phosphate dehydrogenase (GAPDH) connect the important branch point metabolite 3-phosphoglycerate with the upper part of glycolysis and the CBB cycle; the reactions carry high flux during mixotrophic growth, but not during photoheterotrophic growth (Figure S4) (Nakajima et al. 2014; You, He, and Tang 2015). However, we found that the fitness penalty for one of the GAPDH isoenzymes, Gap2, was highest during photoheterotrophic growth while Gap1 and Pgk repression had no penalty (Figure 3 A). Of the two GAPDH isoenzymes in *Synechocystis, gap2* (*sll1342*) has been intensively studied (Valverde, Losada, and Serrano 1997), including its redox-dependent association with the small regulatory protein CP12 (*ssl3364*) and Prk (Michelet et al. 2013; McFarlane et al. 2019; Lucius et al. 2022), and is known to participate in both gluconeogenesis and glycolysis (Valverde, Losada, and Serrano 1997). In contrast, little is known of the physiological role of Gap1 (*slr0884*). Repression of Gap1 showed no fitness effect in any tested condition (Figure 3 A), which is in contrast to a previous study in which a Gap1 mutant could not grow in a mixotrophic condition (Koksharova et al. 1998). We also found that repression of the regulatory protein CP12 resulted in a similar growth pattern to Gap2: significantly reduced growth in mixotrophic and photoheterotrophic conditions. This suggests that CP12 plays a role in regulating the Calvin cycle in the presence of glucose, as recently reported (Blanc-Garin et al. 2022; Lucius et al. 2022). To validate the CRISPRi library results, we constructed three in-frame knockout mutants for *gap1, gap2*, and *cp12* (Figure 3 B-D). We could not obtain a fully segregated Δ*gap2* strain, while Δ*gap1* and Δ*cp12* knockouts were possible. The Δ*gap1*, Δ*gap2* (partial), and Δ*cp12* strains had phenotypes similar to the repression clones from the CRISPR library; Gap1 knockout did not affect growth in photoautotrophic or mixotrophic conditions, while Gap2 and CP12 knockouts reduced growth in the presence of glucose (Figure 3 B-D, Figure S6). It has been shown that the Gap2 of *Synechococcus* can accept NAD(H) or NADP(H) cofactors, but this preference shifts towards NAD(H) upon binding to CP12, and further when the ternary Gap2-CP12-Prk complex is formed, as NADP(H) activity is reduced to zero (McFarlane et al. 2019). Binding of Gap2 and CP12 (and potentially Prk) in dark conditions may thus regulate the change in metabolic flux from gluconeogenesis to glycolysis direction. Our results are thus further evidence for the importance of a fine-tuned regulation on Gap2 for both inorganic carbon uptake by the Calvin cycle and organic carbon assimilation by glycolysis.

### Fitness effects of ncRNAs across growth conditions

The CRISPRi repression library also contains 4950 mutants with sgRNAs targeting 1868 non-coding RNAs (ncRNAs). Of these, 85% were alternative transcriptional units directly associated with a gene such as antisense RNAs (asRNAs) or internal transcription start sites (iTSS) (Table 2). Small RNAs (sRNAs) were 15% of the targeted ncRNAs. sRNAs are presumably independent transcriptional units located between annotated ORFs, and several have been implicated in regulating gene expression (Mitschke et al. 2011; Kopf and Hess 2015). Only a few ncRNAs showed an effect on fitness (Table 2). Of the different ncRNA classes, iTSSs showed the highest proportion of elements with significant effect on fitness and asRNAs the lowest (Table 2, Figure S7 A). This result is not surprising, sgRNAs targeting iTSS will also repress transcription of the coding gene, but with lower efficiency. To evaluate how much of the fitness effect of asRNAs and iTSSs repression was actually caused by repression of the associated genes, we selected all ncRNAs with a significant effect on fitness in at least one condition and correlated this fitness with gene fitness. The correlation of fitness scores between a gene and its corresponding asRNA was high (R = 0.67) while correlation of fitness between a gene and iTSS was lower (R = 0.37) (Figure S7 B,C). Only a few ncRNAs were found that affected cell fitness independent of their associated gene. We therefore focused our analysis on sRNAs and found 27 sRNAs with a strong effect in at least one growth condition (Figure 4 A).

**Table 2:** Summary of ncRNA repression effect in *Synechocystis*. Significance was defined as exceeding a combined score of effect size (fitness) times negative log10 p-value (Wilcoxon rank sum test). Number of significant ncRNAs was determined by counting all ncRNAs that had a combined score ≥ 4 in at least one out of eleven growth conditions.

| ncRNA type | No. of targets | No. of targets with<br>sign. fitness effect<br>(score $\geq 4$) | % sign. targets |
| --- | --- | --- | --- |
| asRNA | 912 | 18 | 1.97 |
| iTSS | 529 | 61 | 11.53 |
| sRNA | 245 | 27 | 11.02 |

**Figure 4.**
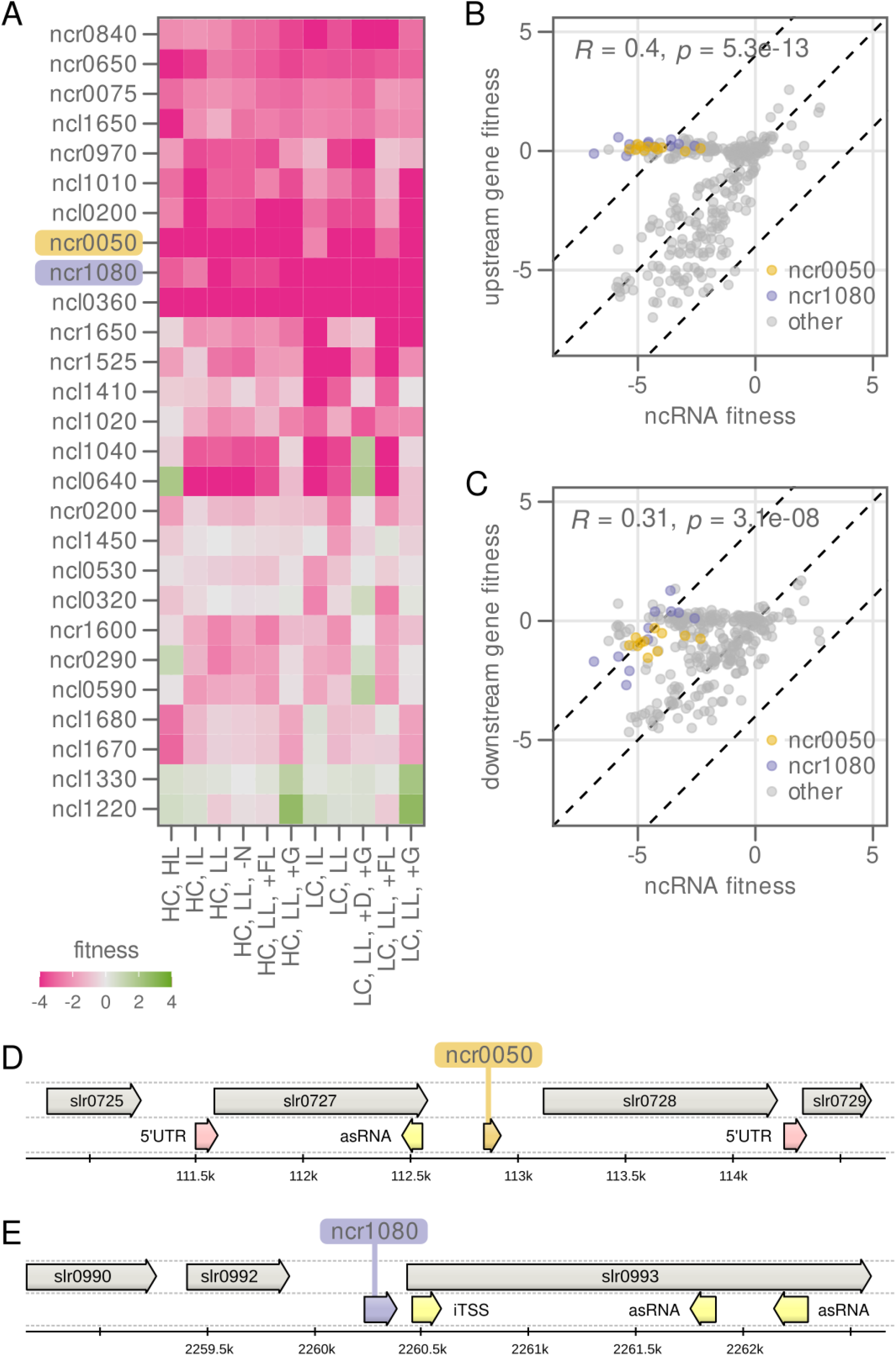
Most ncRNAs do not have an independent effect on fitness. **A)** Heat map showing fitness score of selected small RNAs (sRNAs) in 11 different growth conditions. **B)** Correlation of fitness score of the 27 sRNAs in A) with fitness score of their respective upstream located genes. Every dot represents one sRNA in one growth condition. **C)** As in B) but for the downstream located genes. Colored dots highlight ncRNAs with position independent fitness effect. **D)** Genomic context of the sRNA *ncr0050*. **E)** Genomic context of the sRNA *ncr1080*.

Of the identified sRNAs, many were important for growth in all conditions (*i*.*e*. repression of the sRNA reduced growth rate) while some (e.g. *ncl0530* and *ncl0320*) had more condition-dependent effects (Figure 4 A). Interpretation of the fitness scores of these sRNAs is complicated by possible polarity effects, as repression of an sRNA may also affect expression of the surrounding genes. We therefore compared the fitness score of each sRNA to the fitness score of the co-localized genes (upstream and downstream) in each condition (Figure 4 B,C). This comparison revealed only two ncRNAs that had fitness scores independent of those of their surrounding genes, *ncr0050* and *ncr1080* (Figure 4 D,E). These two sRNAs may be *in trans* to their target, and are important for growth in all tested conditions. Ncr1080 (also named SyR47), is located upstream of the lipoprotein nlpD (*slr0993*) but forms an independent transcriptional unit that is upregulated in high light (Kopf et al. 2014). Ncr0050 is not previously characterized.

### Prediction of sgRNA efficacy from sequence

Most genes in the *Synechocystis* CRISPRi library were targeted by five sgRNAs. This redundancy made it possible to correlate fitness scores (a measure of sgRNA efficacy) of each sgRNA with its nucleotide sequence. It was shown previously that the efficacy of the guide RNA in mediating DNA cleavage by the Cas9 nuclease varies, and that some of this variation can be attributed to the binding properties of the RNA/DNA hybrid (Xu et al. 2015; Labuhn et al. 2018; Peng et al. 2018; Xiang et al. 2021). Consequently, numerous algorithms have been derived in these and similar studies in order to predict the efficacy of guide RNAs (Liu, Zhang, and Zhang 2020), most recently using deep learning approaches (L. Wang and Zhang 2019; Xiang et al. 2021). Most of these studies focused on the CRISPR/Cas9 DNA cleavage system and eukaryotic hosts, while prokaryotes and CRISPRi repression systems each have their own sequence requirements (L. Wang and Zhang 2019; Xu et al. 2015). Here, we leveraged the information obtained in our extensive fitness screening to predict sequence motifs that lead to better or worse repression. To this end, sgRNA sequences including 8 to 12 nt of the 5’ and 3’ genomic context were selected as features for statistical learning, together with five additional features derived from the sgRNA as a whole (GC content, melting temperature, length, distance to promoter, and ‘crisproff’ score) (Alkan et al. 2018). Features were used to train an ensemble of four different models (random forest, gradient boosting machine, support vector machine, multi-layer perceptron, Figure 5 A). The models were trained to predict a binary classifier based on the repression efficiency *E* (low: 0 ≤ *E* ≤ 0.5, high: 0.5 < *E* ≤ 1). From the complete set of gene-targeting sgRNAs (n=16,476), we included only those where the targeted gene had any effect on fitness in at least one condition, even if the effect was small (abs(fitness) > 1, n=6,306, Figure 5 B). This comparatively low fitness cutoff was chosen to include a sufficient number of sgRNAs for training. Of these sgRNAs, 3,841 were labeled as low-efficiency and 2,465 as high-efficiency sgRNAs. This dataset was further split into a training and a validation set containing 75% and 25% of the data, respectively. Models were built and trained using the python packages scikit-learn and keras/tensorflow (Methods) and model quality was evaluated by comparing the predicted sgRNA class with the actual class in the validation set. In terms of sensitivity (ability to retrieve high-efficacy sgRNAs), the best performing model was the support vector machine followed by the multi-layer perceptron, while decision tree-based models had problems retrieving high-efficacy sgRNAs (Figure S8 A, Table S1). Of the 606 high-efficacy sgRNAs in the validation set, 386 were correctly labeled by at least one out of four models (Figure 5 B, C), while 220 were not correctly labeled by any model. Overall model performance was nevertheless far from perfect: on average only 41% of the high-efficacy sgRNAs were correctly identified. Several problems might contribute to difficulties in identifying a certain subset of high-efficacy sgRNAs: The type or number of features is too limited; The number of observations (sgRNAs) is too low for the models in order to pick up important patterns during training; Unknown or random regulatory events in the cell influence sgRNA efficacy. Next, we were wondering which features were most important for the models to classify sgRNAs. The two decision tree based models (random forest, gradient boosting machine) record feature importance, here meaning the influence of every base at every position in the sgRNA as well as its direct neighborhood (Figure 5 C, Figure S9). Positions -4 to -12 were of particular importance, more specifically a cytosine residue (C) at position -4, a stretch of guanine residues (G) from position -5 to -12, and a stretch of thymidine residues (T) from position -7 to -10. Inspecting the sequence motifs of the different sgRNA classes using logo plots revealed a clear enrichment of sequence patterns (Figure 5 D). While a pool of all used sgRNAs showed only a slight preference for G/C for the entire length of the sequence, high-efficiency sgRNAs that were identified by at least one model (n=386) showed a strong enrichment of G and a depletion of T in position -5 to -11, in accordance with feature importance. Interestingly, high-efficiency sgRNAs that were not identified showed a less specific pattern, explaining the difficulty for models to retrieve them correctly. From the five additional features calculated from the entire sequence, three were contributing to sgRNA classification: the distance to the promoter was the most important feature, followed by ‘crisproff’ score and melting temperature (Figure S8 B). Interestingly, sgRNA length and GC content had little importance, although the previously mentioned G-stretch was a prominent sequence motif. We next validated that GC content was not an important trait differentiating high-and low-efficacy sgRNAs (Figure S8 C). The two groups had only a marginal difference in average GC content (55% and 53%, respectively). The difference between correctly identified and unidentified high-efficacy sgRNAs was higher though (57% and 51% respectively). We conclude that G enrichment and T depletion in the ‘seed region’ (first 12 bases preceding the PAM site) is a good predictor for sgRNA efficiency, although not the only one.

**Figure 5.**
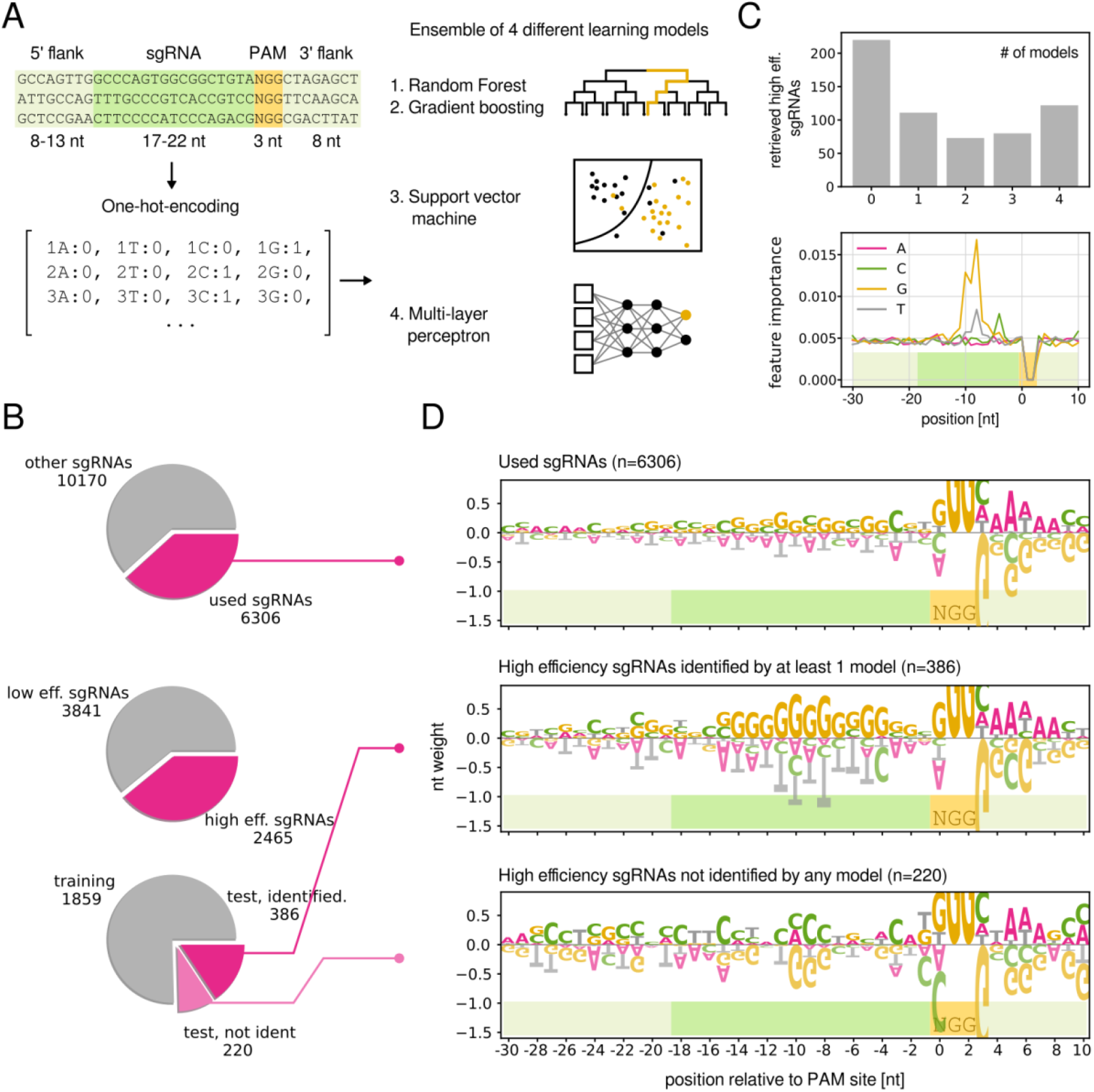
Prediction of sgRNA efficiency from sequence. **A)** Schematic overview of model training and prediction. SgRNA sequences were one-hot-encoded, split into training and validation sets, and used to train four different learning models. **B)** Subsets of sgRNAs used for model training and testing. Of the 6,306 sgRNAs included in modeling, 3,841 were labeled as high efficiency and 2,465 as low efficiency. Of the 606 high efficiency sgRNAs in the validation (test) set, 386 were labeled correctly by at least one model. **C)** Number of correctly labeled sgRNAs in the test set by number of models (upper panel). Feature importance in nucleotide species per sequence position derived from random forest (lower panel). See Figure S9 for the equivalent feature importance derived from gradient boosting. **D)** Sequence logos visualizing the weight of each nucleotide at each position (Tareen and Kinney 2020). Height of letters corresponds to the probability of encountering the respective nucleotide. Positive values, enrichment, negative values, depletion.

## Discussion

By performing growth competition cultivations across 11 conditions, we were able to identify condition-specific essentiality of many genes, including those with unknown function (Figure S1, Supplemental_material_2). These genes would be a high priority for future study for the cyanobacterial community. Comparison across conditions also revealed unexpected fitness contributions of known genes, such as the importance of Gap2 and CP12 in glycolysis metabolism, and the relative unimportance of Gap1 (Figge et al. 2000) (Figure 3). This finding is in contrast to a previous, widely cited study reporting that a Gap1 mutant completely lost the capacity to metabolize glucose (Koksharova et al. 1998). However, an earlier study had shown that Gap2 is the only enzyme from *Synechocystis* that compensates for GAPDH function in *E. coli* (Valverde, Losada, and Serrano 1997). Furthermore, the *Synechocystis* Gap1 is phylogenetically related to the plant cytosolic GapC, does not show GAPDH activity *in vitro*, and is expressed at a low level (Valverde, Losada, and Serrano 1997; Jahn et al. 2018). While Gap2 repression negatively affected growth in mixotrophic and photoheterotrophic conditions, we did not observe a growth penalty upon Gap2 repression or partial knockout (a full knockout was not possible) in phototrophic conditions, which may be due to the fact that in both cases, a base amount of Gap2 was retained that was sufficient to support growth. Therefore, as the ancestor of plant GAPDH (Martin and Cerff 2017), cyanobacterial Gap2 may possess both chloroplastic NADPH-dependent GAPDH activity and cytosolic NADH-dependent GAPDH function, while the existence of Gap1, which is thought to be an NADH-dependent GAPDH, may be an evolutionary ancestor of the widespread second GAPDH in plants. The PSII subunit PsbK also showed an anomalous pattern in fitness data; it was the only photosystem gene that showed a strong negative fitness effect upon repression even in photoheterotrophic and mixotrophic conditions. Therefore, while other PSII subunits appear to be dispensable when glucose is present (Ikeuchi et al, 1990), PsbK may have a universal role in electron transport which is not restricted to PSII function.

The suboptimal regulations we observed were mainly for proteins within light harvesting and conversion: phycobilisomes, PSII subunits, and Flv2-4 proteins. The improved growth upon repression of PBS could be due to reduced electron pressure in the ‘HC,HL’ and photoheterotrophy condition, as even quenched PBS still allows 60% of absorbed energy to reach the reaction centers (Kirilovsky 2015). Antennae knockout was also shown to increase PSII and decrease PSI content in *Synechocystis* (Luimstra et al. 2019). An alternative hypothesis is that repression of PBS provides a benefit on protein economy, as predicted by metabolic modeling (Jahn et al. 2018). ApcAB and CpcAB are the most abundant PBS proteins (Figure S3), so their repression would free the most proteome space for additional ribosomes to fuel growth in the ‘HC, HL’ condition where cells grew fastest. Furthermore, repression of CpcC, a linker protein of low abundance but important for antenna light harvesting (Lea-Smith et al., 2014), did not show a growth increase. However, repression of ApcE, a protein that may anchor Apc discs to the thylakoid and is much smaller than ApcAB (Domínguez-Martín et al. 2022), showed similar, if weaker, fitness profiles as ApcAB. The similarity suggests that a reduced antennae connectivity, and not necessarily protein burden may also contribute to the growth rate increase, though it is not clear to what extent ApcAB still forms in the ApcE mutant; both ApcAB and ApcE knockouts were described as ‘olive’ mutants with completely abolished PBS function (Ajlani et al. 1995). The growth advantage of PSII subunits and PBS in the photoheterotrophic condition (and an overall weaker effect in the mixotrophic condition) point to the repression of functional light harvesting and conversion as being beneficial. Flv2-4 catalyze an alternative electron flow pathway in cyanobacteria and the observed growth advantage upon repression of Flv2-4 suggests that they are excessively expressed in our cultivation conditions. In typical air-lift photobioreactors (PBRs) for microalgae, insufficient mixing can lead to dark zones so that cells experience larger fluctuations in light on the order of seconds, even in constant illumination (Andersson et al. 2019). In these reactors, alternative electron flow was shown to be highly active and likely beneficial (Andersson et al. 2019)). However, our turbidostat cultures used air-lift PBRs with short path length and low cell density, so that light fluctuations due to mixing were mitigated and Flv2-4 proteins became a burden. Suboptimal regulation of antennae and alternative electron flow in cyanobacteria is reminiscent of sustained NPQ in plants, which are typically slow in adapting to stress due to low expression of recovery enzymes or other factors (Malnoë 2018). For example, energy dependent quenching (qE) in plants is beneficial for light fluctuations, but sustained qE after the fluctuation has ended hinders photosynthesis (Kaiser, Morales, and Harbinson 2018; Kromdijk et al. 2016). While photoprotection mechanisms in plants are more diverse than in cyanobacteria, there are some parallels, such as the recently discovered slowly relaxing photoprotective quenching (qH) which may have a homologue in cyanobacteria (Gorbunov et al. 2011; Staleva et al. 2015; Amstutz et al. 2020). The use of a CRISPRi sgRNA library in the background of cyanobacteria strains deficient in known photoprotection mechanisms could identify additional players.

CRISPRi allows for controlled gene repression, which is beneficial for studying the role of genes across conditions: consider that a knockout of a gene essential for photoautotrophic growth would remove that clone from the population already after transformation. However, a complication is that gene repression is not total; residual enzyme or protein may be retained so that a phenotype is not observed. For example, the repression of PsaK2 (*sll0629*) did not show a fitness change in any condition (Figure S2 A), even though PsaK2 was identified as an essential element involved in state-transition based non-photochemical quenching during acclimation to high light (Fujimori, Hihara, and Sonoike 2005). Under high light conditions, expression of PsaK2 is highly elevated to assist energy transfer from phycobilisome to PSI. This upregulation may result in an excess amount of protein to ensure cell robustness in light-stress conditions, and CRISPRi repression mutants may retain a basal level of PsaK2 that is sufficient to avoid a growth penalty. Another possible example of insufficient gene repression by dCas9 is of Flv2-4 in low C_i_ conditions (Figure 2 C). *Synechocystis* strongly upregulates Flv2-4 expression in low C_i_ conditions, due to their role as an acceptor of excess electrons downstream of PSI. Flv2-4 knockout mutants showed impaired photosynthesis (Shimakawa et al. 2015; Santana-Sanchez et al. 2019). However, in the CRISPRi library, we did not observe a negative effect of Flv2-4 repression in all LC conditions, rather the opposite. Thus, we expect that even reduced Flv2-4 amounts are sufficient for cells to cope with light fluctuations in these reactors.

This work improves significantly upon a previous CRISPRi library (Yao et al. 2020), both in terms of growth conditions screened and in library quality. The inclusion of five sgRNAs targeting each gene was critical for a confident assessment of gene fitness. The binding efficiency of sgRNAs is highly sequence dependent, and off-target binding is a significant challenge. For example, an analysis of dCas9 binding in *E. coli* found that sgRNAs with PAM regions containing a 9-nt identity elsewhere were likely to bind off-target (Cui et al. 2018). Therefore, the effect of all sgRNA clones for a target gene should be considered when determining fitness scores. We found that the multiple hypothesis adjusted p-value (padj) associated with each fitness score was a good filter to prevent false conclusions from single outlier sgRNAs with strong fitness effects. In cases where the fitness score was associated with a low p_adj_ (high significance), the fitness effect could be reproduced by gene-knockout (*e*.*g*. Figure 3B-D), but not in cases where p_adj_ was high (low significance, Figure S10). Because the gene repression by CRISPRi is on the transcriptional level, the fitness score of neighboring, co-transcribed genes must also be considered when interpreting the fitness score of a gene of interest.

The retrospective analysis of sgRNA efficacy using our comprehensive fitness data revealed properties important for sgRNA design in cyanobacteria. Firstly, sgRNA design principles for dCas9 mediated CRISPR interference need to be different from principles used for Cas9 mediated DNA cleavage. Similar to the study from (Xu et al. 2015; Labuhn et al. 2018; Peng et al. 2018; Xiang et al. 2021), we found that a region in the center of the spacer (−4 to -12) is most important for sgRNA efficacy, rather than the four most proximal positions to the PAM site which were deemed important for Cas9-mediated cleavage (Xu et al. 2015; Labuhn et al. 2018; Peng et al. 2018; Xiang et al. 2021; D. Wang et al. 2019). The importance of the central region was reported previously but only in conjunction with high GC content (Labuhn et al. 2018). Here, the region showed a marked preference for G but not C, and in-depth analysis of sequence features confirmed that GC content alone is not an important variable. The genomic context surrounding the spacer played no role in determining guide efficacy. From the higher-order features, distance to the promoter was most important (Xu et al. 2015; Labuhn et al. 2018; Peng et al. 2018; Xiang et al. 2021) followed by the ‘crisproff’ score which estimates the binding energy of the DNA-sgRNA hybrid (Alkan et al. 2018). The effect of promoter distance was stronger in our library than in a comparable pooled CRISPRi library for *E. coli* (M. N. Price et al. 2018; T. Wang et al. 2018; Jahn et al. 2021; Yao et al. 2020; Garst et al. 2017; Vo et al. 2021). For prediction of sgRNA efficacy from plain sequence, distance to the promoter might not always be available for scientists. If it is known, we recommend to select sgRNAs which target the first 100 nt downstream of the start codon where efficacy is highest. The promoter itself or the 5’ UTR was not targeted in this study although other studies successfully targeted these elements.

## Methods

### sgRNA library design

Up to 5 sgRNAs were designed for each open reading frame (ORF) and ncRNA. ORF were retrieved from NCBI (reference genome assembly ASM972v1, accessed on 18.03.2016) and ncRNA locations were obtained from Kopf and Hess (Kopf and Hess 2015) as previously described (Yao et al. 2020). An in-house Python script available at [https://github.com/KiyanShabestary/library_designer] was used to design protospacer sequences using the following criteria: GC content between 40% and 80%, absence of bad seeds (Vigouroux and Bikard 2020), absence of G_6_ and T_4_, length between 18 and 23bp. Target sequences were searched with the pattern 5’-[CCN]-(N_18_-N_23_)-3’ on the coding strand (NGG PAM). Off-targets were screened on both forward and reverse strands for sites containing either the canonical NGG or the alternative NAG PAMs. Guiding RNAs with homologies at most one mutation away in the 15 bp adjacent to the PAM were discarded. For a given ORF or ncRNA, guiding RNAs that were at least 5 bp from one another were considered.

### Genetic construction of sgRNA library

The library was inserted in a *Synechocystis* base strains containing a tetR_PL22_dCas9/SpR expression cassette genome integrated at the psbA1 locus. The sgRNA oligos were synthesized on two 12 K chips (Custom Array inc., USA), pooled together in equimolar ratio. The sgRNA oligos were then cloned into an entry vector targeting the slr0397 locus using Golden-gate assembly, as previously described (Yao et al. 2020). The ligation mix was transformed into NEB 10-beta competent E. coli cells and approximately 1’000’000 colonies were obtained. Colonies were collected in LB, pooled and grown over-night. Plasmid DNA was extracted using the ThermoFisher Maxi plasmid extraction kit. Plasmid (10µg) was transformed in *Synechocystis* tetR_PL22_dCas9/SpR base strain via natural transformation. After 10 days at 30°C and constant illumination, colonies were collected in BG-11 and pooled. The pooled library was stored in 7% DMSO at -80°C.

### Turbidostat cultivation in photobioreactors

The *Synechocystis* sgRNA library was cultivated in 8-tube Multi-Cultivator MC-1000-OD bioreactors (Photon System Instruments, Drasov, CZ) with 65 mL culture volume per tube. Temperature (30°C), constant light intensity (from back side of the photobioreactor), and turbidostat pumping system were controlled by an in-house computer program described in Jahn et al., 2018. A gas mixing system GMS150 (Photon System Instruments, Drasov, CZ) was used to provide 1% v/v CO_2_ for HC conditions and air otherwise. Gas bubbling rate was manually set to an average of 90 bubbles per minute, with a total flow rate of 100 mL/min per reactor tube. Fluctuating light (1500 µmol photons m^−2^ s^−1^) was provided by an extra LED light panel, PARADIGM LIGHT WH 1200-V (Beambio), from the front side of the photobioreactor. Culture OD_720 nm_ and OD_680 nm_ were automatically measured every 15 min by the photobioreactor, and the turbidity threshold was set to OD_720 nm_ = 0.2. Once the threshold was exceeded for 3 measurements in any tube, 5 mL fresh medium was pumped into the tube for dilution. All cultures were initially cultivated in turbidostat at a standard condition (30°C, 60 µmol photons m^−2^ s^−1^, BG11 (pH=7.8) medium with 25 µg mL^-1^ spectinomycin, 25 µg mL^-1^ kanamycin, and 0.5 µg mL^-1^ anhydrotetracycline) for 48 hours to allow sufficient repression, after that specific conditions (Table 1) were applied. Generation time was calculated as T_gen_ = ln 2/growth rate, and T_0_ was the condition switching time point. At 4^th^, 8^th^, and 10^th^ generation, cells were harvested by centrifuging 12 mL culture at 4°C, 3000 × g for 10 min. Supernatant was discarded completely and cell pellets were stored at −20°C.

### Library preparation and next-generation sequencing

Genomic DNA was extracted from harvested cell pellets using GeneJET Genomic DNA purification Kit (Thermo Fisher Scientific), using the protocol for Gram-positive bacteria due to the dense cyanobacterial cell wall. Extracted gDNA was used as template for the 1^st^ PCR to amplify sgRNA region and add NGS adaptors. PCR products were purified using AMPure XP beads (BECKMAN COULTER) and used as template for the 2^nd^ PCR where Illumina barcodes were added by NEBNext Multiplex Oligos for Illumina (Dual Index Primers Set 1 and 2) (New England Biolabs). PCR products were purified using AMPure XP beads (BECKMAN COULTER), and quantified by the Qubit™ 4 Fluorometer (Thermo Fisher Scientific). Samples were pooled together such that the final concentration was equal (100 ng/µL), and the pooled library was purified from agarose gel using GeneJET Gel Extraction Kit (Thermo Fisher Scientific). Two rounds of NGS were carried out on an Illumina NextSeq 2000 system using NextSeq 2000 P3 kit (50 Cycles), with 72 samples sequenced simultaneously per round.

### Fitness score calculation

Sequencing data was preprocessed using an automated pipeline available at https://github.com/m-jahn/nf-core-crispriscreen. Sequencing data was retrieved from Illumina’s base space server using the bs-cp tool. Fastq.gz files were quality trimmed using Trimgalore v0.6.7 (https://www.bioinformatics.babraham.ac.uk/projects/trim_galore/). Read and run quality was summarized using FastQC v0.11.9 (https://www.bioinformatics.babraham.ac.uk/projects/fastqc/) and MultiQC v1.12 (https://multiqc.info/). Reads from filtered fastq.gz files were aligned to the reference genome (fasta file with all sgRNA sequences) using Bowtie2 v2.4.4 (Langmead et al. 2019) with option -U (unpaired reads). The output from Bowtie2, sequence alignment files (SAM files), were further processed by samtools (Danecek et al. 2021) using view and sort commands. Counts of mapped reads per sgRNA were calculated using subread/featureCounts v2.0.1 (https://nf-co.re/modules/subread_featurecounts). Next, count tables per sample were summarized to a single table using a custom R script, and statistics for pairwise sample comparison were calculated using DESeq2 (Love, Huber, and Anders 2014). Log_2_ fold change values were normalized between conditions using the function normalize Between Arrays (method=“quantile”) from the R package limma to account for possible differences in number of generations. Fitness scores for each sgRNA and condition were calculated by determining the area under the curve of the log_2_ fold change (*log*_*2*_ *FC*) over time *t* normalized by total cultivation time.

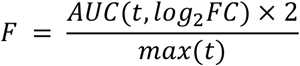

In the second step, a single fitness score for each gene was calculated from all sgRNAs of that gene by determining the weighted mean, where weight *wi* for each sgRNA *i* was based on the correlation coefficient *Ri* of one sgRNA with the others, and its repression efficiency *E*_*i*_

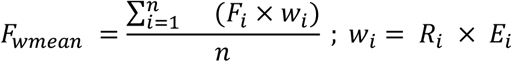

The repression efficiency *E* is the fitness score of a single sgRNA divided by the maximum fitness of all sgRNAs for the same gene (0 ≤ *E* ≤ 1).

### Statistical analysis

All cultivation and sequencing experiments were performed with three biological replicates. Replication was carried out at the stage of bioreactor cultivation (inocula were obtained from single pre-cultures grown in 200 mL shake flasks). P-values were calculated from a comparison of sgRNA fitness score (n = 1 to 5 depending on gene) with non-targeting control sgRNAs (n=10) using the Wilcoxon rank sum test. P-values were multiple-hypothesis corrected using the Benjamini-Hochberg procedure. For selected tasks, a combined score *S* was calculated for each target and condition by combining effect size (fitness score F) and adjusted p-value according to the formula:

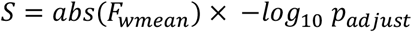

A combined score threshold of 4 corresponds to an absolute fitness score of 2 and an adjusted p-value of 0.01 and was considered as significant. All analyses of fitness data were performed using the R programming language and are documented in an R notebook available at https://github.com/m-jahn/R-notebook-crispri-lib.

### Construction and cultivation of single knockout mutants

Since Gap1 (*slr0084*), Gap2 (*sll1342*), CP12 (*ssl3364*), and *slr1505* are individually located in the *Synechocystis* genome, chloramphenicol cassette was integrated into corresponding gene locus to generate the knockout strains. Integrating plasmids were designed to have 1000 bp homologous regions on both upstream and downstream of the target gene. Genotypes of the knockout strains were confirmed using two pairs of primers, one pair anneal to chloramphenicol cassette to screen colonies, and another pair anneal to the original gene to check segregation. The knockout strains were cultivated in batch mode with initial OD730 = 0.1 in multicultivators. The specific conditions used for the batch cultivation were kept identical to the corresponding conditions in turbidostat cultivation.

### Machine learning to predict sgRNA efficiency

The machine learning pipeline was implemented as a jupyter notebook available at https://github.com/m-jahn/R-notebook-crispri-lib, using python v3.9.7. In order to predict sgRNA efficiency, sgRNA sequences were imported from a fasta file and mapped back to the genome to retrieve the PAM site and 5’ and 3’ genomic flanks. Flanking sequences were variable in length to allow a fixed total length of 40 nt. Nucleotide sequence was one-hot-encoded by converting an A, T, C, or G at a single position into a 4-digit binary vector of 0 or 1. Altogether this resulted in a vector of 160 features for each sgRNA. Only a subset of 6,306 sgRNAs was used for modeling, where the targeted gene showed a fitness score of abs(F) ≥ 1 in at least one condition. The target variable for training was the sgRNA efficiency E, ranging between 0 and 1, and experimentally determined from CRISPRi library cultivations. To simplify modeling, E was binned into two categories, ‘low’ (E < 0.5) and ‘high’ (E ≥ 0.5). An ensemble of four models for classification problems were selected, RandomForestClassifier, GradientBoostingClassifier, and support vector machine (svm) from scikit-learn, as well as a muli-layer perceptron (small, fully connected neural network) from keras/tensorflow. The feature array and the target variable were split into training and validation data sets (75% or 4730 observations and 25% or 1576 observations, respectively). The first three models were tuned using a grid search with the following parameters. Random forest: ‘n_estimators’: [20, 50, 100, 200, 300, 400], ‘max_features’: [‘auto’, ‘sqrt’], ‘max_depth’: [5, 10, 25, 50, 100]. Gradient boosting: ‘loss’: [‘deviance’, ‘exponential’], ‘n_estimators’: [100, 200, 300], ‘learning_rate’: [0.01, 0.05, 0.1, 0.5], ‘max_depth’: [1]. Support vector machine: ‘kernel’: [‘rbf’, ‘linear’], ‘C’: [0.5, 0.75, 1, 1.5, 2], ‘gamma’: [0.01, 0.05, 0.1, 0.5, 1, 5]. Model training was then performed with the best parameter set. For the multi-layer perceptron, different configurations of 1 to 3 hidden layers with 8 to 128 nodes were tested manually, and the best performing configuration selected. The final topology of the model was: 1 input layer with 64 nodes and activation function ‘relu’, 1 hidden layer with 128 nodes and activation function ‘relu’, and 1 output layer with 2 nodes and activation function ‘softmax’. The model was trained with maximally 200 epochs but already converged after less than 100. The loss function that evaluates model performance during training was ‘categorical_crossentropy’, and the optimizer was gradient descent with momentum (‘tf.keras.optimizers.experimental.SGD’) with learning rate = 0.01 and momentum = 0.9.

## Data and software availability

All data and data processing pipelines are publicly available on the following platforms: Raw sequencing data was deposited at the European Nucleotide Archive (ENA accession number XXX). Sequencing data was processed using a Nextflow data analysis pipeline which is available on https://github.com/MPUSP/nf-core-crispriscreen. Analysis of processed data with R 4.1.2 is documented in R markdown notebooks available at https://github.com/m-jahn/R-notebook-crispri-lib. All used software is open source and publicly available on web-accessible resources. Fitness data can be browsed interactively using the R Shiny app available at https://m-jahn.shinyapps.io/ShinyLib/.

## Acknowledgment

We thank Ute Hoffmann for helpful discussions on ncRNA results. We also thank Emil Sporre and Markus Janasch for assistance with bioreactor cultivation. This study was funded by the Novo Nordisk Foundation NNF20OC0061469, Swedish Research Council 2020-04329, Swedish Foundation for Strategic Research SSF ARC19-0051, and the Swedish Research Council Formas (Grant number 2015-939 and 2019-01491).

## Author contributions

R.M. performed cloning of strains, cultivation experiments, library preparation, NGS sequencing, designed the study, and wrote the manuscript. M.J. performed data analysis, statistical analysis, machine learning, designed the study, and wrote the manuscript. K.S. designed the guide RNA library and transformed the library into *Synechocystis*. E.P.H. designed the study and wrote the manuscript. All authors read and approved the final manuscript. All authors agreed that the position of authors who contributed equally may be swapped for personal applications, curricula vitae, or similar purposes.

