## Supplemental_material_1 for "CRISPR interference screens reveal tradeoffs between growth rate and robustness in *Synechocystis* sp. PCC 6803 across trophic conditions"

for the the manuscript entitled

CRISPRi, CRISPR interference, gene repression, high throughput, functional genomics, gene function prediction, cyanobacteria, photoautotrophy, stress resistance, robustness, growth rate, fitness, *Synechocystis*

**Table S1**. Performance metrics of machine learning models. SVM - support vector machine, GBM - gradient boosting machine, RF - random forest, MLP - multi-layer perceptron. Two different classes were predicted, low-efficacy sgRNAs ("0-low") and high-efficacy sgRNAs ("1-high"). * - accuracy is identical for both classes because it is calculated taking both classes into account.

| **model** | **class** | **precision** | **recall / sensitivity** | **accuracy*** | **f1_score** |
|---|---|---|---|---|---|
| SVM | 0-low | 0.684 | 0.710 | 0.619 | 0.696 |
| SVM | 1-high | 0.504 | 0.474 | 0.619 | 0.489 |
| GBM | 0-low | 0.675 | 0.850 | 0.656 | 0.752 |
| GBM | 1-high | 0.589 | 0.345 | 0.656 | 0.435 |
| RF | 0-low | 0.669 | 0.792 | 0.630 | 0.725 |
| RF | 1-high | 0.527 | 0.371 | 0.630 | 0.436 |
| MLP | 0-low | 0.651 | 0.658 | 0.573 | 0.655 |
| MLP | 1-high | 0.443 | 0.436 | 0.573 | 0.439 |

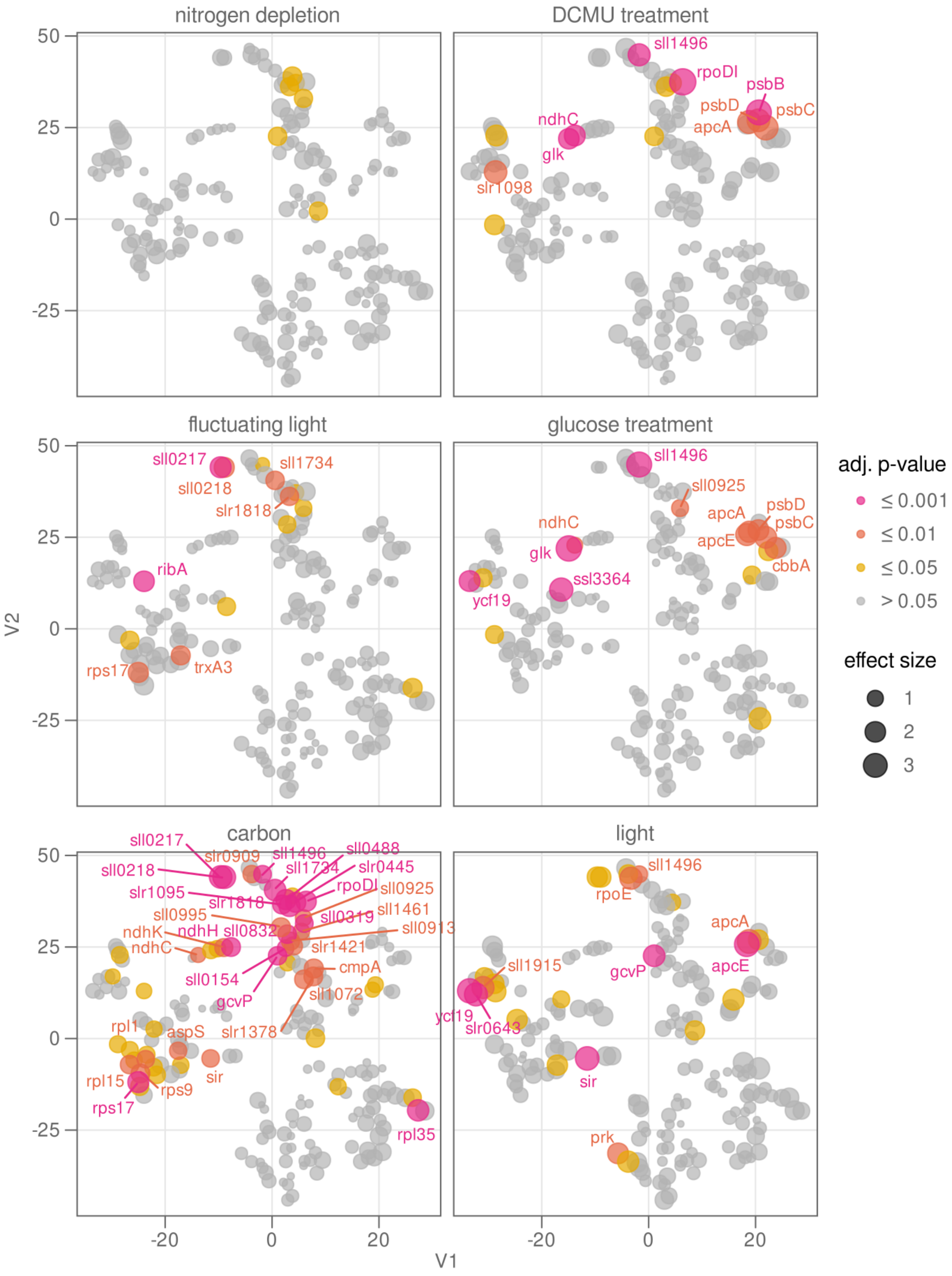


**Figure S1.** Bubble plot showing t-SNE clustering of 187 genes which were selected based on absolute fitness score ≥ 4 and adjusted p-value ≤ 0.01 (Wilcoxon rank sum test). Bubble size encodes effect size of the respective treatment (carbon, light, additional treatments) from multiple linear regression models. Bubble color encodes adjusted p-value from Wilcoxon rank sum test. V1, V2, two dimensions showing gene similarity.

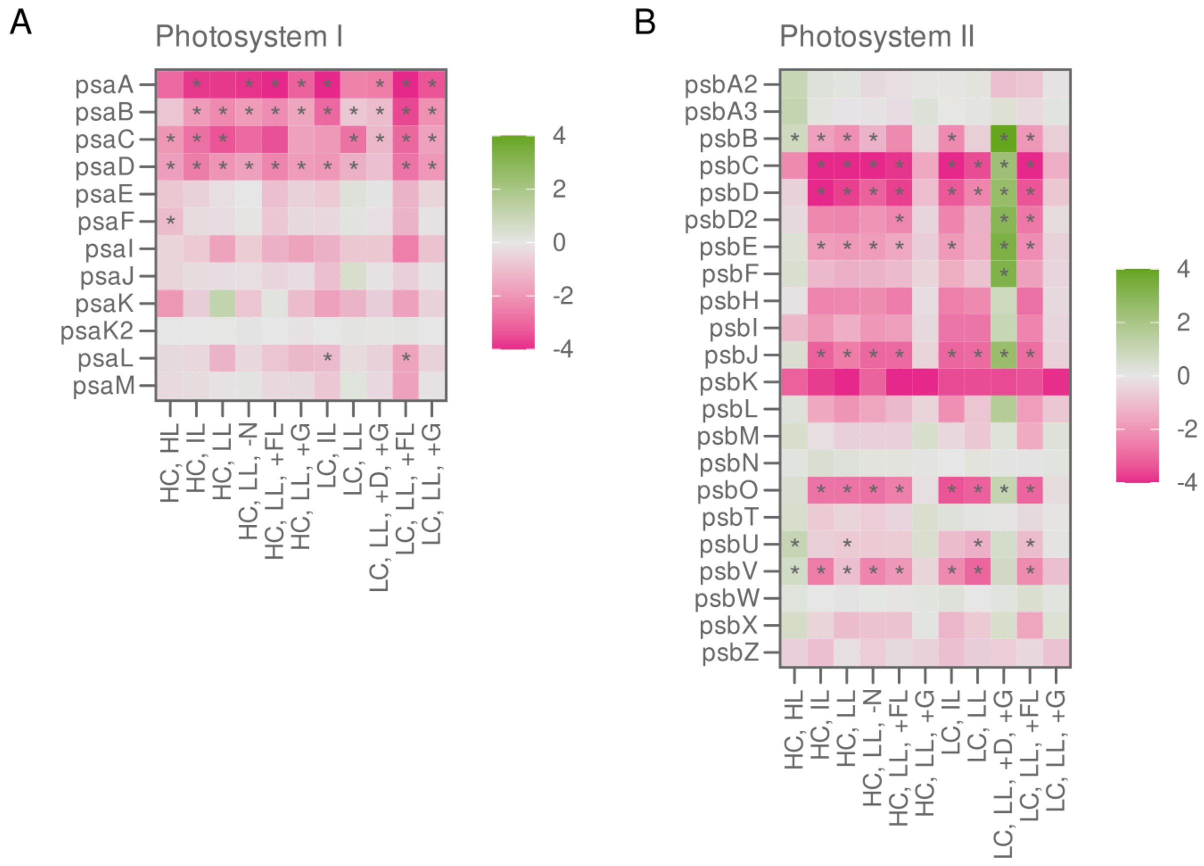


**Figure S2.** Heatmap showing fitness score for repression of photosystem genes in *Synechocystis* PCC 6803. **A)** Genes encoding photosystem I subunits. **B)** Genes encoding photosystem II subunits. Asterisk: Wilcoxon rank sum test, adjusted p-value ≤ 0.01.

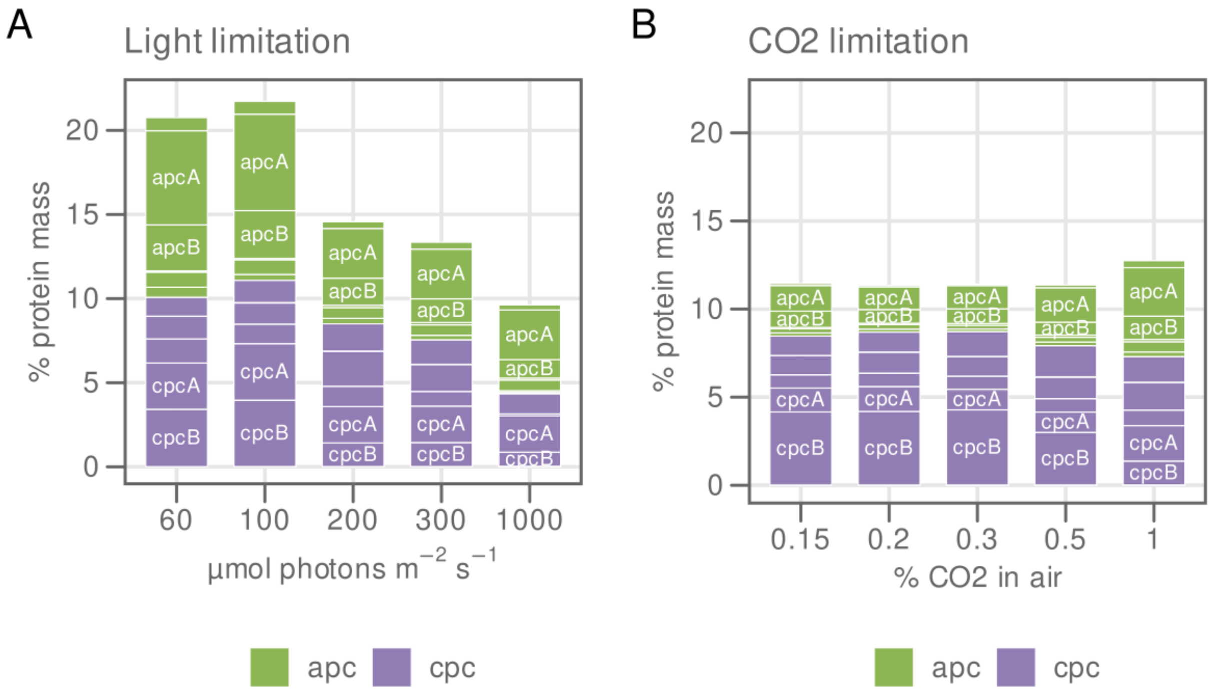


**Figure S3.** Percent protein mass of the total proteome for the light harvesting complex (phycobilisomes) from Synechocystis PCC 6803. Protein mass fraction was estimated from label-free quantification of mass spectrometry data (Jahn et al. 2018). Green, allophycocyanin subunits. Purple, phycocyanin subunits. **A)** Protein mass depending on light limitation. **B)** Protein mass depending on carbon limitation.

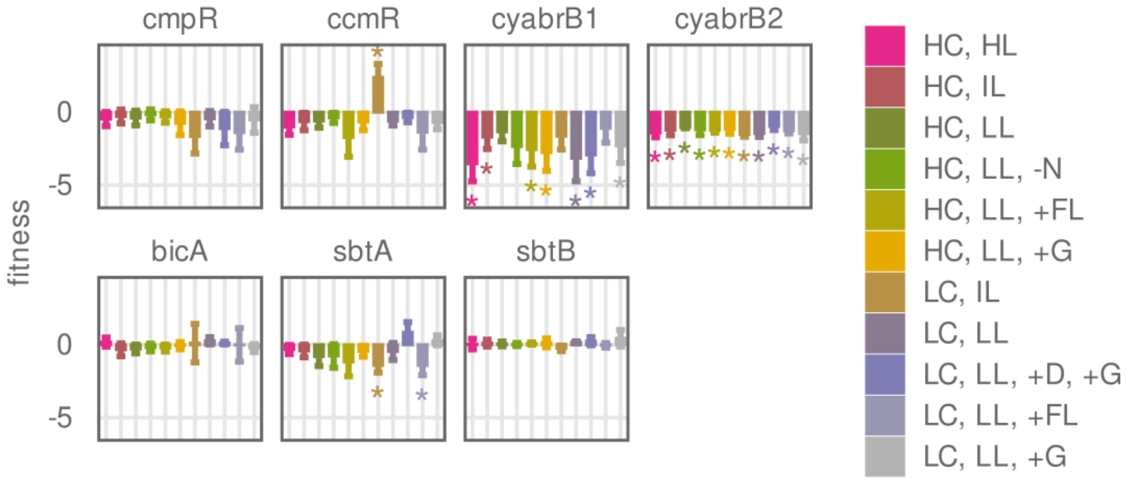


**Figure S4.** Fitness score for repression of genes involved in carbon transport. *CmpR*, *ccmR*, *cyabrB1*, and *cyabrB2* are regulatory genes. *BicA*, *sbtA* and *sbtB* are carbon transporters. Asterisk: Wilcoxon rank sum test p-value ≤ 0.01.

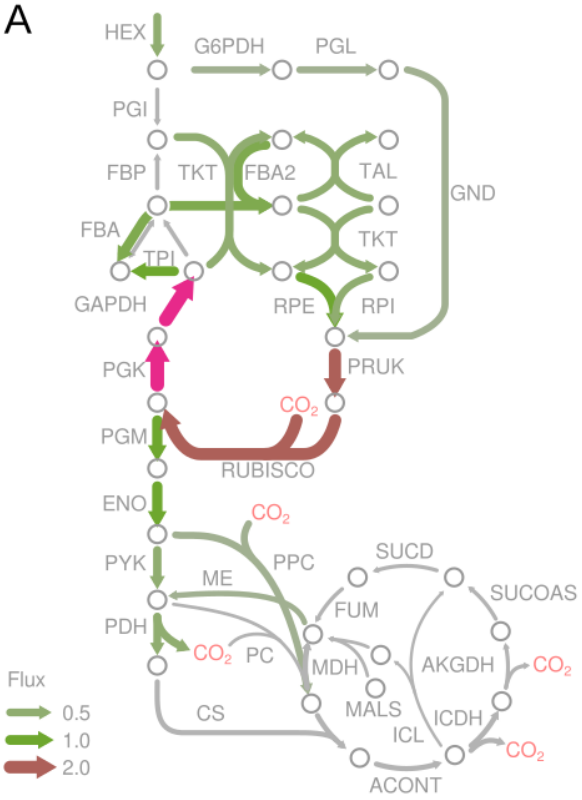


**Figure S5.** Metabolic flux for mixotrophic and photoheterotrophic growth of *Synechocystis* sp. PCC 6803. Fluxes were determined by (Nakajima et al. 2014) using $^{13}C$ labeled D-glucose. Fluxes were plotted on a custom made metabolic map using the R package fluctuator (*https://github.com/m-jahn/fluctuator*). Reactions and their directionality are shown with arrows and named with capital letters according to the BiGG standard. Arrow thickness indicates the reaction flux in mmol g $DCW^{-1}$ $h^{-1}$. **A)** Mixotrophy (growth in presence of glucose). **B)** Photoheterotrophy (growth in presence of glucose and DCMU to block photosystem activity).

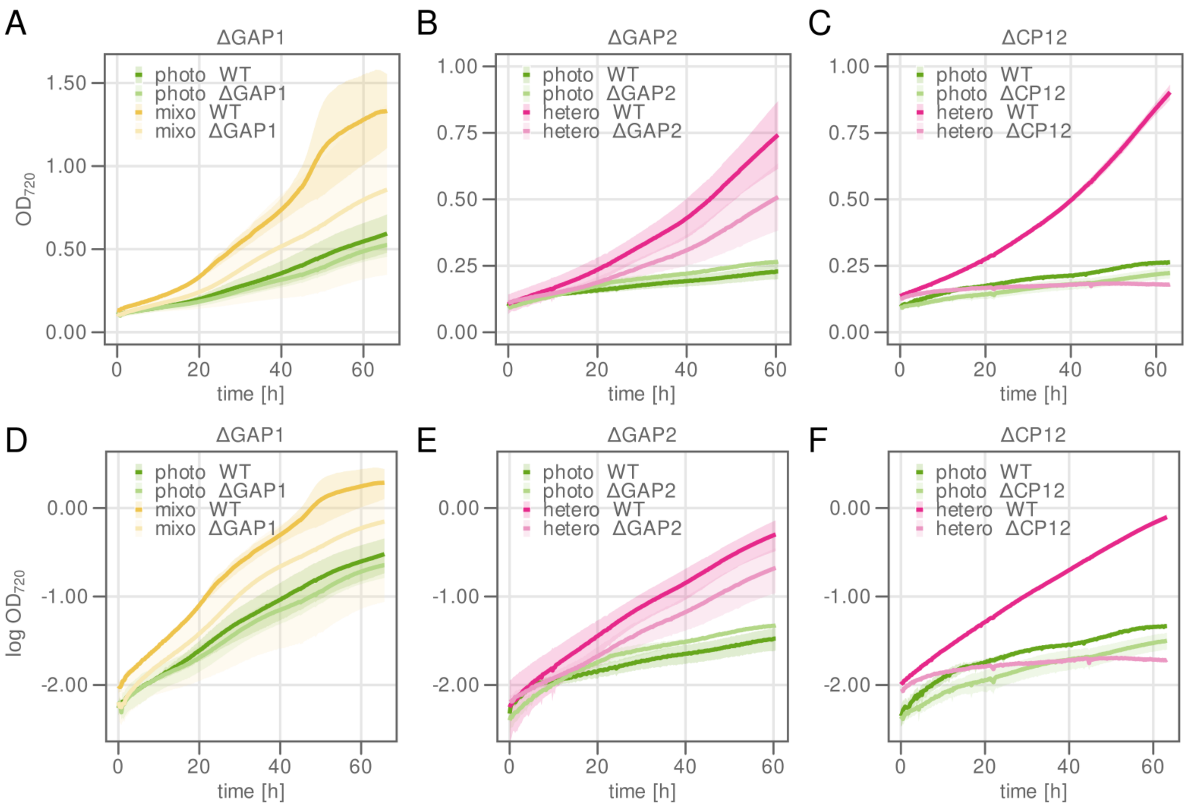


**Figure S6.** Validation of CRISPRi library results for *gap1*, *gap2* and *cp12* using deletion mutants. Cultivation was performed in the same conditions as were used for cultivating the CRISPRi library, but in batch mode instead of turbidostat. WT was used as control in each cultivation. Growth recorded as change in optical density at 720 nm ($OD_{720}$). Line and ribbons: Mean and standard deviation of at least two biological replicates. **A)** Δ*Gap1* mutant. Phototrophy: HC, LL. Mixotrophy: HC, LL, +G. **B)** As in A) but for Δ*gap2*. Phototrophy: LC, lL. Photoheterotrophy: LC, LL, +G, +D. **C)** As in A) but for Δ*cp12*. Phototrophy: LC, LL. Photoheterotrophy: LC, LL, +G, +D. **D-F)** As in A-C) but natural logarithm-transformed.

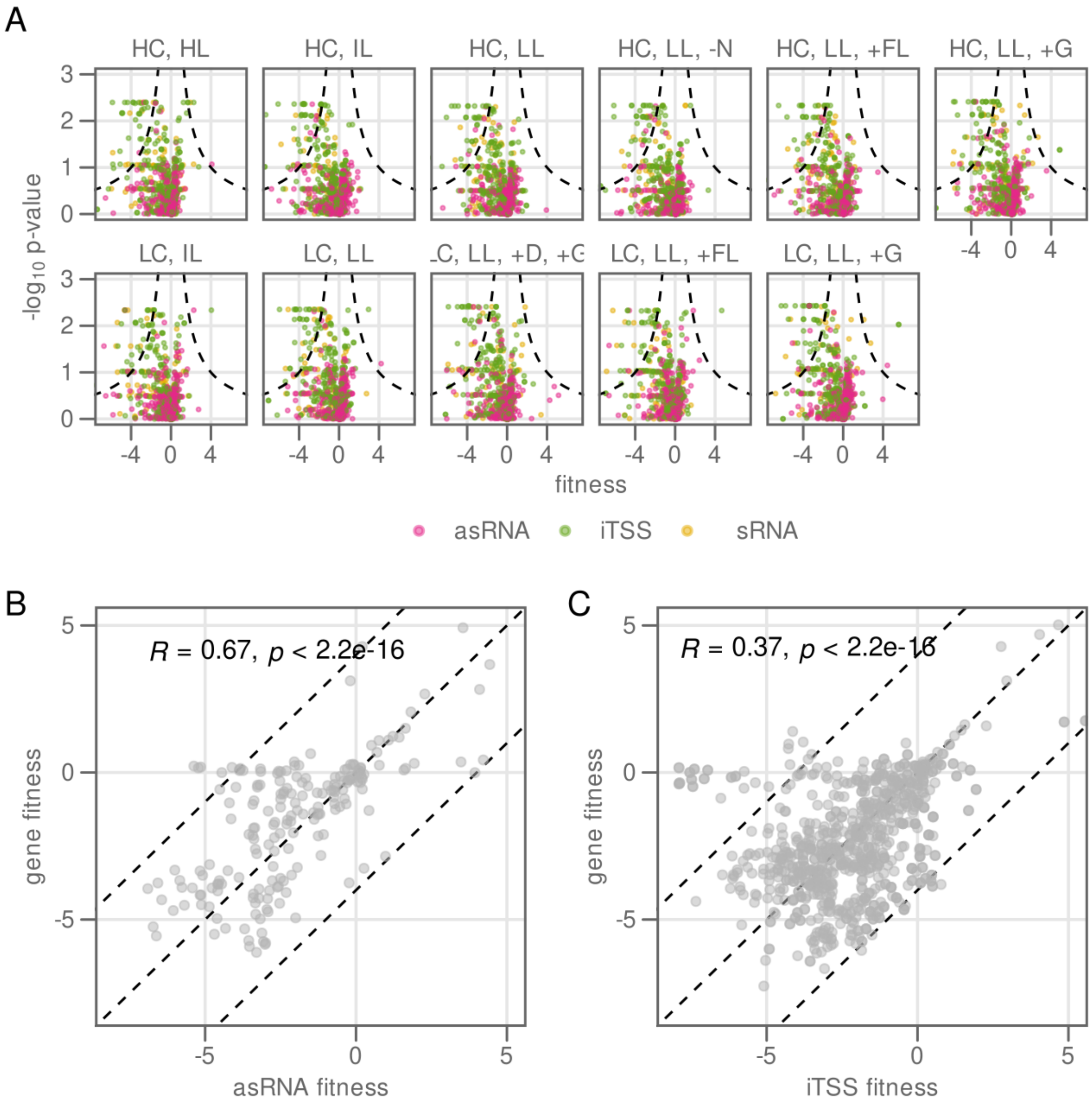


**Figure S7.** Non-coding RNAs (ncRNAs) targeted by the CRISPRi library. **A)** Volcano plot showing fitness (x-axis) and negative $\log_{10}$ p-value from Wilcoxon rank sum test (y-axis) for all ncRNAs, broken down by growth condition and type. asRNA, antisense RNA. iTSS, internal transcription start site. sRNA, small RNA. **B)** Correlation of fitness score of all asRNAs with the fitness score of their respective sense-oriented gene. Every dot represents one asRNA in one growth condition. *R*, correlation coefficient. Dashed lines: boundaries for an absolute fitness difference of 4 between gene and asRNA. **C)** As in B) but for the iTSSs.

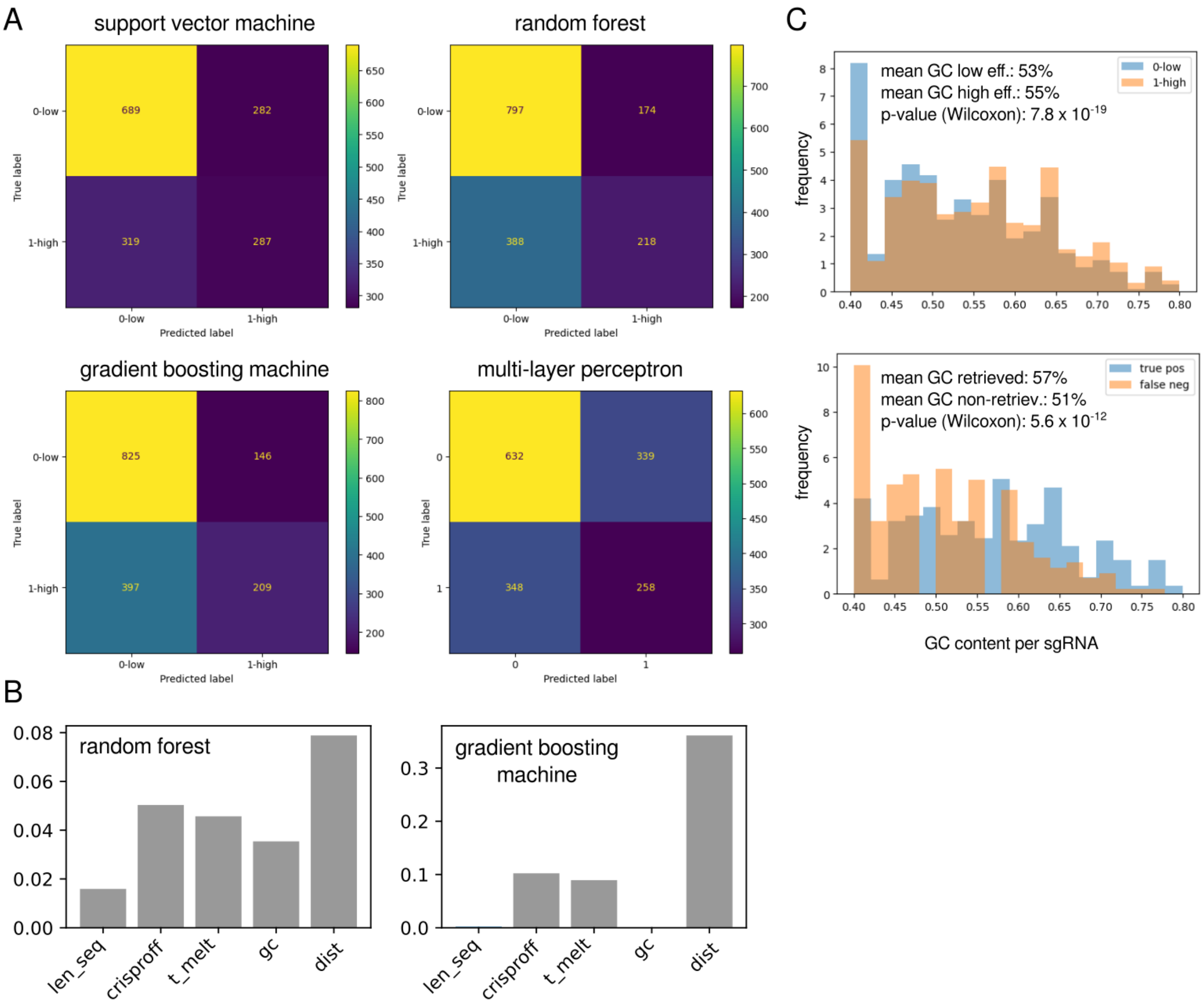


**Figure S8.** Additional statistics for sgRNA efficacy machine learning. **A)** Confusion matrices for all four used models showing number of correctly and incorrectly retrieved sgRNA classes. Upper left quadrant: true negatives, upper right: false positives, lower left: false negatives, lower right: true positives. **B)** Feature importance for five additional features derived from sequence or genomic context. len_seq: length of sgRNA (17-22), crisproff: on-target score calculated using the CrisprOff tool at https://github.com/RTH-tools/crisproff (Alkan et al. 2018), t_melt: melting temperature of sgRNA, gc: GC content, dist: distance to the promoter. **C)** Detailed analysis of GC content for different groups of sgRNAs. Upper panel: Histogram of GC content between low efficacy and high efficacy sgRNAs. Lower panel: Histogram of GC content for correctly retrieved (true positive) and incorrectly retrieved (false negative) high efficacy sgRNAs.

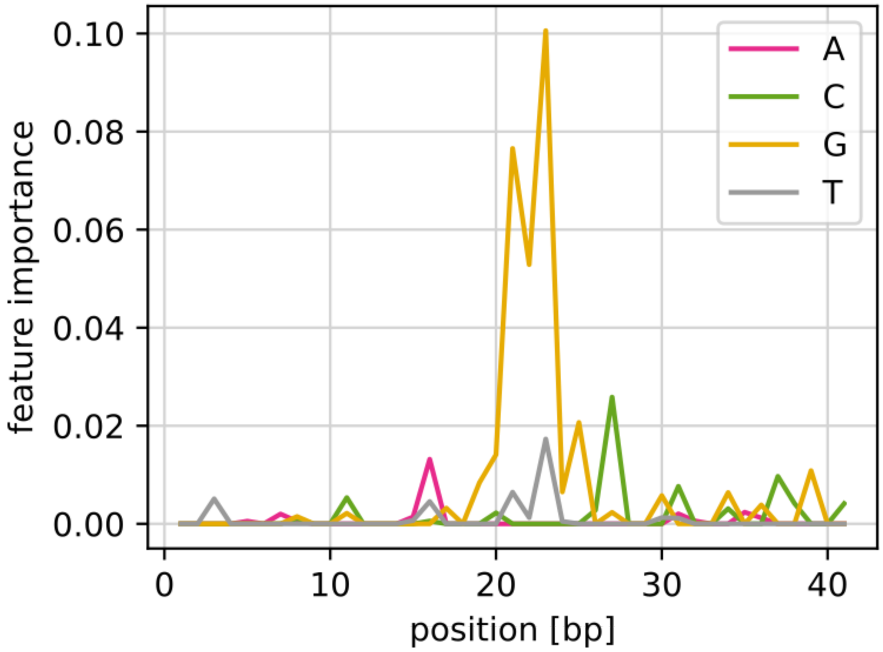


**Figure S9.** Feature importance in nucleotide species per sequence position derived from gradient boosting model. See Figure 5 C in the main text for the equivalent feature importance derived from the corresponding random forest model.

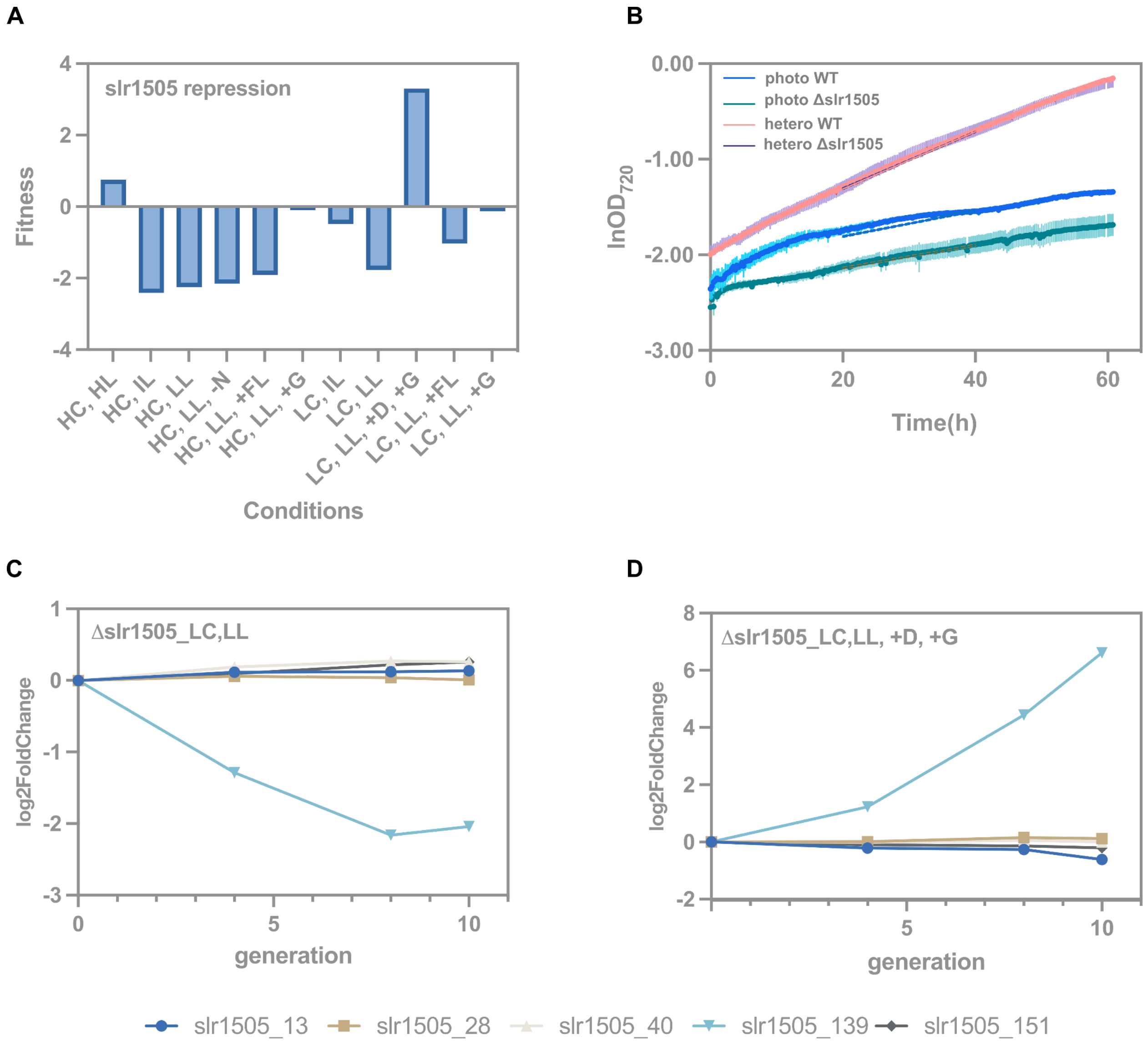


**Figure S10.** Validation example of gene with high absolute fitness score but high *p*-value. **A)** Fitness scores of slr1505 in all 11 conditions tested on CRISRPi library. **B)** Natural logarithm-transformed $OD_{720}$ of WT and Δ*slr1505* in 60 hours batch cultivation. **C)** Log2FoldChange along 10 generations of 5 individual sgRNAs target slr1505 in phototrophic condition. **D)** Log2FoldChange along 10 generations of 5 individual sgRNAs target slr1505 in photoheterotrophic condition.
